# Functional *in vivo* characterization of zebrafish *sox10* enhancers in melanoma and neural crest

**DOI:** 10.1101/2020.02.06.932178

**Authors:** Rebecca L. Cunningham, Eva T. Kramer, Sophia K. DeGeorgia, Shayana Seneviratne, Vadim Grigura, Charles K. Kaufman

## Abstract

The re-emergence of a neural crest transcriptional program, including Sox10 upregulation, is a key step in melanoma initiation in humans and zebrafish. We hypothesize that epigenetic regulation of *sox10* modulates melanoma initiation. ATAC-Seq analysis of zebrafish melanoma tumors identifies recurrently open chromatin domains near *sox10*. Reporter constructs for each putative *sox10* enhancer were examined in zebrafish embryos for neural crest activity and in stable transgenic lines for melanoma activity. One element, *peak5* (23 kilobases upstream of *sox10*), drives *EGFP* reporter expression in a subset of neural crest cells, Kolmer-Agduhr neurons, and early melanoma patches and tumors with high specificity. A ∼200 bp region, conserved across the *Cyprinidae* family (fish), is required for *peak5* activity in neural crest and melanoma, and contains dimeric SoxE binding sites essential for neural crest activity. Our work identifies a novel melanoma transcriptional enhancer, expanding our knowledge of epigenetic regulation of neural crest identity in melanoma.

## Introduction

Melanoma is a potentially deadly cancer of transformed melanocytes, which are pigment-producing cells derived from embryonic neural crest. Early detection and treatment of melanoma is critical because of high mortality associated with metastatic melanoma. The difficulty of treating metastatic melanoma underscores the necessity of defining how this cancer initiates and the urgency of developing new therapeutics (Lo and Fisher, 2014). Encouragingly, chromatin modifiers and other proteins that shape the epigenetic landscape offer promising novel therapeutic targets that may expand the treatment repertoire beyond immunotherapy and MAPK-targeting strategies (Tanaka et al., 2015; Badner et al., 2017), and combat the ability of melanomas to evade treatment due to the high mutational burden, heterogeneous mutations, and rapid evolution of these tumors (Gallagher et al., 2014; Fontanals-Cirera et al., 2017).

To effectively treat melanoma in its earliest stages requires a thorough understanding of mechanisms that drive melanoma initiation. The transgenic *BRAF*^*V600E*^*/p53*^*-/-*^ zebrafish has been established as a powerful melanoma model that is highly reflective of the human disease and can be utilized to study early stages of melanoma initiation (Patton et al., 2005). In this model, the human *BRAF*^*V600E*^ oncogene is expressed under the control of the melanocyte-specific promoter of the *mitfa* gene. Within *BRAF*^*V600E*^*/p53*^*-/-*^ zebrafish, all melanocytes thus have the potential to become melanoma, but only one to three melanomas typically develop from the many thousands of melanocytes in any single fish, presenting an excellent model in which to study how cancer initiates from a field of cancer-prone cells.

Recent work in zebrafish has demonstrated that reactivation of a neural crest program (NCP) is an early step in melanoma initiation (Kaufman et al., 2016). A single melanoma cell can be visualized in live zebrafish by expression of the *crestin:EGFP* transgene (Kaufman et al., 2016). Not only does *crestin:EGFP* expression uniquely enable visualization of cancer before a large tumor is visible, expression of *crestin* also indicates a re-emergence of aspects of neural crest transcriptional identity in these cells because *crestin* is exclusively expressed in the embryonic neural crest of zebrafish (Luo et al., 2001). Neural crest cells (NCCs) are a transient and migratory embryonic cell population that give rise to a variety of cell types, including melanocytes. Prior studies have noted that NCCs and cancer cells share many cell biological characteristics, including the ability to proliferate, change cell adhesion properties, and migrate (Maguire et al., 2015). For example, epithelial-to-mesenchymal transition occurs to permit both NCC migration and cancer metastasis (Kerosuo and Bronner-Fraser, 2012) Furthermore, melanoma cells can behave as NCCs after transplantation into chick embryos, demonstrating their plasticity and connection to their developmental origins (Kulesa et al., 2006).

While *crestin* is expressed in the neural crest, this gene is zebrafish-specific and functionally uncharacterized; therefore, *crestin* itself does not represent a future therapeutic target in human melanoma. However, *crestin* is one of a suite of neural crest genes that are upregulated in melanoma in zebrafish and humans (Kaufman et al., 2016). The re-emergence of an NCP transcriptional program in melanoma is supported by the expression of other neural crest related genes in tumors such as *sox10* and *dlx2a*. Therefore, applying knowledge of developmental mechanisms in NCCs may enhance our understanding of melanoma initiation and treatment of NCC derived cancers.

One highly upregulated neural crest gene in melanoma in both zebrafish and humans is *sox10*, a transcription factor necessary for neural crest development (Kaufman *et* al., 2016; Rönnstrand and Phung, 2013; Kelsh, 2006). Furthermore, in a zebrafish *BRAF*^*V600E*^ melanoma model, a *Nras* mouse melanoma model, and in human melanoma cell culture, abrogation of *sox10* expression affects melanoma onset and maintenance, respectively (Kaufman *et al*., 2016; Shakhova *et al*., 2012). Likewise, overexpression of *sox10* in melanocytes increases the rate of melanoma onset in zebrafish (Kaufman *et al*., 2016). Further demonstrating an NCP activation in melanoma, regulatory elements near neural crest genes, such as *sox10*, are activated in melanoma cells lines and NCCs, but not generally in other adult tissues or selected cancer cells lines (Kaufman *et al*., 2016). Importantly, this epigenetic phenomenon is conserved in human melanomas (Kaufman *et al*., 2016).

Since *sox10* is critical to establish neural crest identity, the development of melanocytes, and the maintenance and progression of melanoma (Kaufman *et al*., 2016; Shakhova et al., 2012; Cronin et al., 2013), epigenetic regulation of *sox10* may be critical for re-establishing neural crest identity in melanoma initiation. Therefore, identifying and functionally characterizing enhancer elements that control *sox10* expression during this oncogenic transition will broaden our understanding of epigenetic changes that occur to trigger melanoma initiation and activation of an NCP-like identity from within a cancerized field.

In this study, we functionally screen putative enhancers elements, identified as accessible or “open” regions of chromatin using ATAC-Seq on zebrafish melanomas, near the *sox10* gene for transcriptional activating activity in embryonic neural crest as well as in melanoma tumors. This screening approach allowed us to identify and focus initially on one 669 bp enhancer, termed *peak5*, that is highly and specifically active in adult zebrafish melanoma. We further show that *peak5* is an active embryonic enhancer in a subset of the NCCs and a select population of Kolmer-Agduhr neurons in the central nervous system. We also identify a ∼200 bp nucleotide sequence centered within *peak5* that is conserved between members of the fish *Cyprinidae* family and is necessary and sufficient for neural crest activity. Furthermore, mutational analysis of a dimeric SoxE binding site in the conserved core of *peak5* uncovers transcription factor binding sites that are necessary for robust *peak5* neural crest activity. Together, these data enhance our understanding of epigenetic changes that drive the re-emergence of neural crest identity in melanoma and suggest a model wherein auto-regulation of *sox10* in a feed-forward loop may contribute to the formation of melanoma.

## Results

### Identification of putative sox10 enhancers

To identify putative regulatory elements surrounding the *sox10* locus, ATAC-Seq was performed on bulk melanoma tumors isolated from *Tg(BRAF*^*V600E*^*;crestin:EGFP);p53*^*-/-*^ zebrafish (**Figure 1A**). ATAC-Seq on melanoma tumor cells showed consistent regions of open chromatin and identified 11 regions of open chromatin around and within the *sox10* locus, including a putative *sox10* minimal promoter encompassing exon 1 of *sox10* (**Figure 1B; Figure S1**). These open regions of chromatin were also consistent with ATAC-Seq peaks in zebrafish melanoma cell lines (Kaufman et al., 2016) (**Figure S1**). To assess if these ATAC-Seq peaks are functional enhancers in neural crest and melanoma cells, we performed *in vivo* reporter assays in *Tg(BRAF*^*V600E*^*); p53*^*-/-*^ zebrafish embryos (**Figure 1C**). Putative non-intronic enhancers that were present in at least 4 out of 5 tumor ATAC-Seq samples were cloned upstream of the mouse *beta-globin* basal promoter driving *EGFP* (Tamplin et al., 2011), within a Tol2 vector (394 from the Tol2 Kit) to enable mosaic integration into the zebrafish genome.

**Figure 1:**
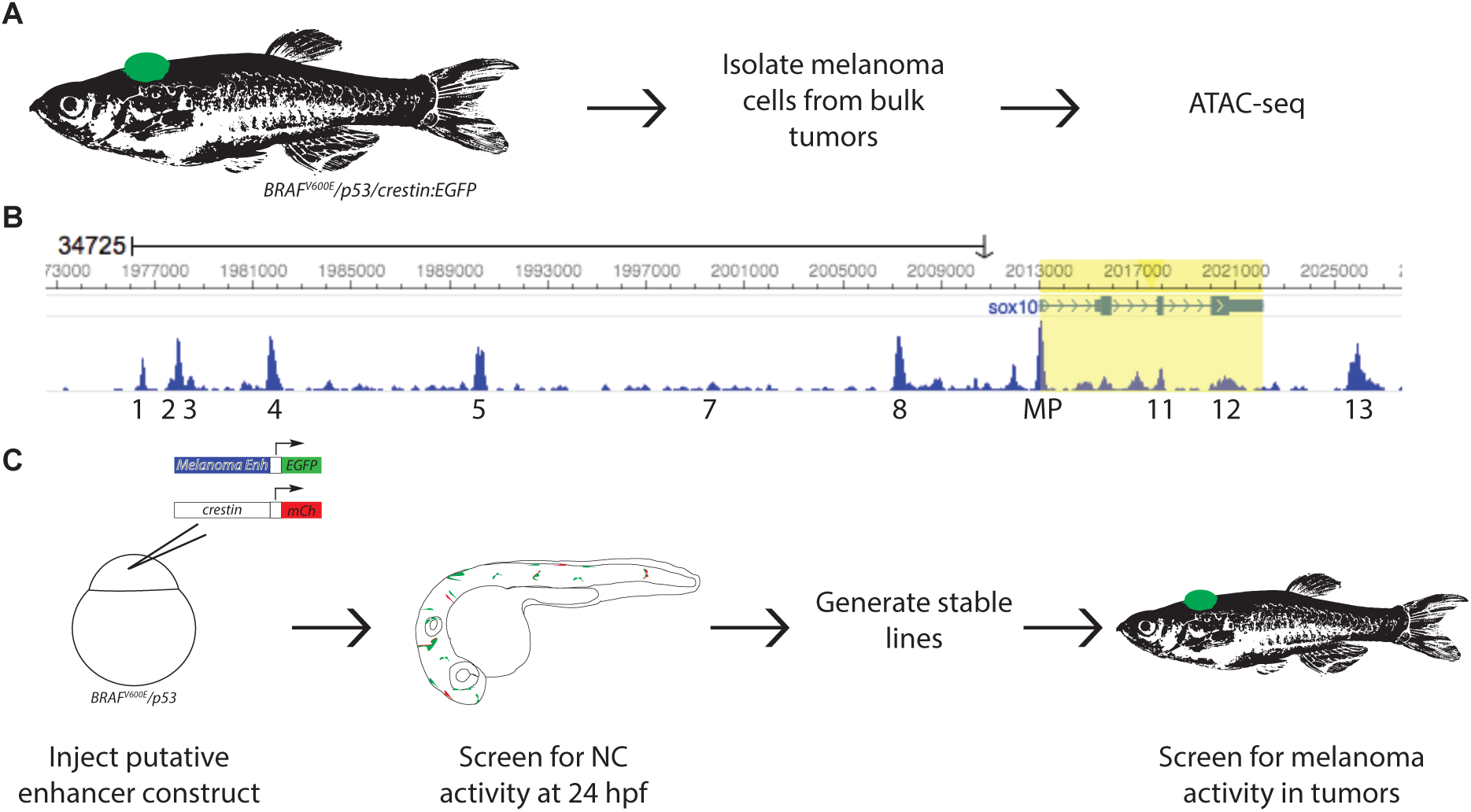
ATAC-Seq identifies putative *sox10* enhancers in zebrafish melanoma tumors. **A)** Schematic for obtaining zebrafish melanoma tumor cells for ATAC-Seq. **B)** Regions of open chromatin surrounding the *sox10* locus. Peak numbers are annotated below. MP = minimal promoter. **C)** Schematic for screening putative enhancers for activity in NCCs and melanoma.

Each putative enhancer construct was injected into single-cell *Tg(BRAF*^*V600E*^*); p53*^*-/-*^ embryos. A *crestin:mCh* construct was also co-injected with each putative enhancer construct to serve as a positive injection control and to co-label NCCs. Embryos were screened at 1 dpf for co-labeling of EGFP and mCh, indicating that the enhancer functions in the neural crest. Larva were ranked as either “no expression,” “weak expression,” or “strong expression” for both *EGFP* and *mCh*. “Strong expression” was characterized by 5 or more EGFP positive cells where as “weak expression” was characterized by fewer than 5 EGFP positive cells. Nine out of eleven putative enhancers drove *EGFP* expression in at least a subset of NCCs, as determined by cell shape, location, and co-labeling with *crestin:mCh* (**Figure S2**). One peak of interest, *peak5*, cloned as a 669 bp element (**Figure S3**), exhibited particularly robust activity in NCCs (**Figure 2A-B**). The putative *sox10* minimal promoter, cloned as a 351 bp element and potential positive control, exhibited the most robust *EGFP* expression throughout the nervous system and in NCCs (**Figure 2C**). A region of closed chromatin approximately 76 kb upstream of *sox10*, as determined by ATAC-Seq, was also cloned upstream of *betaglobin:EGFP* as a negative control and did not drive neural crest *EGFP* expression (**Figure 2D**). In subsequent analyses, we noted that this control region also exhibits sequence similarity to another region on Chromosome 3 (Chr3: 45240516-45241216). An additional negative control (Negative Control A) cloned from a region 10.7 KB upstream of the *sox10* also exhibited minimal reporter activity (**Figure S2**). Collectively, ATAC-Seq analysis of zebrafish melanoma tumors reliably identifies transcriptional regulatory regions near *sox10* that function as enhancers with neural crest cell activity in transgenic reporter studies.

**Figure 2:**
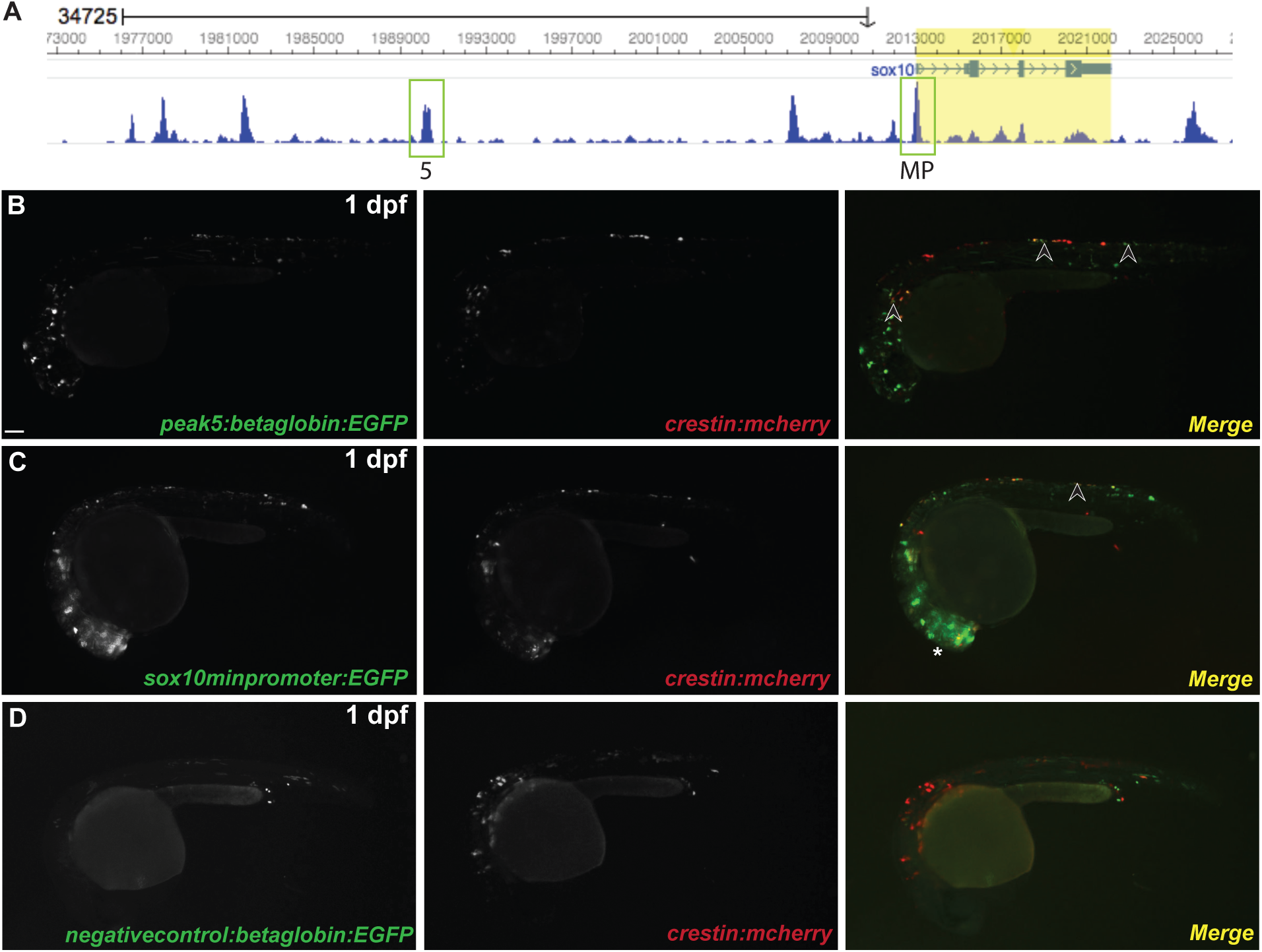
*peak5* is an active neural crest enhancer. **A)** Location of *peak5* and the *sox10* minimal promoter (MP) based on ATAC-Seq. **B)** *peak5-*driven *EGFP* expression is mosaically present in NCCs (arrowheads) in F0 injected embryos, as indicated by co-localization with *crestin:mch* at 1 dpf. **C)** The *sox10* minimal promoter in F0 embryos is active in both NCCs (arrowheads) and the CNS (asterisk). **D)** F0 embryos injected with a negative control (∼76 kb upstream of the *sox10* TSS) exhibit only limited expression of *EGFP* not localized to NCCs.

### peak5 is active in embryonic neural crest and Kolmer-Agduhr neurons

Enhancer reporter constructs injected into zebrafish embryos are randomly integrated throughout the genome, and thus subject to position effects (Udvadia and Linney, 2003). In addition to examining numerous F0 transgenic embryos for consistent spatial and temporal expression as above, we further characterized the expression pattern of activity of *peak5*, a region showing particularly robust neural crest activity in zebrafish embryos, by generating 6 independent stable transgenic lines. To generate stable lines, F0 injected embryos were raised to maturity and then outcrossed to *Tg(BRAF*^*V600E*^*); p53*^*-/-*^ zebrafish. Subsequent F1 embryos were then screened for *EGFP* expression at 1 dpf. All 6 identified stable transgenic lines exhibited commonalities in expression patterns, detailed below (**Figure 3**). One stable line, *peak5_115A* exhibited EGFP signal in muscle. Given that *sox10* is not endogenously expressed in muscle, we reasoned that this muscle activity was due to position effects, such as enhancer trapping, and as a result did not use this line for subsequent analyses.

**Figure 3:**
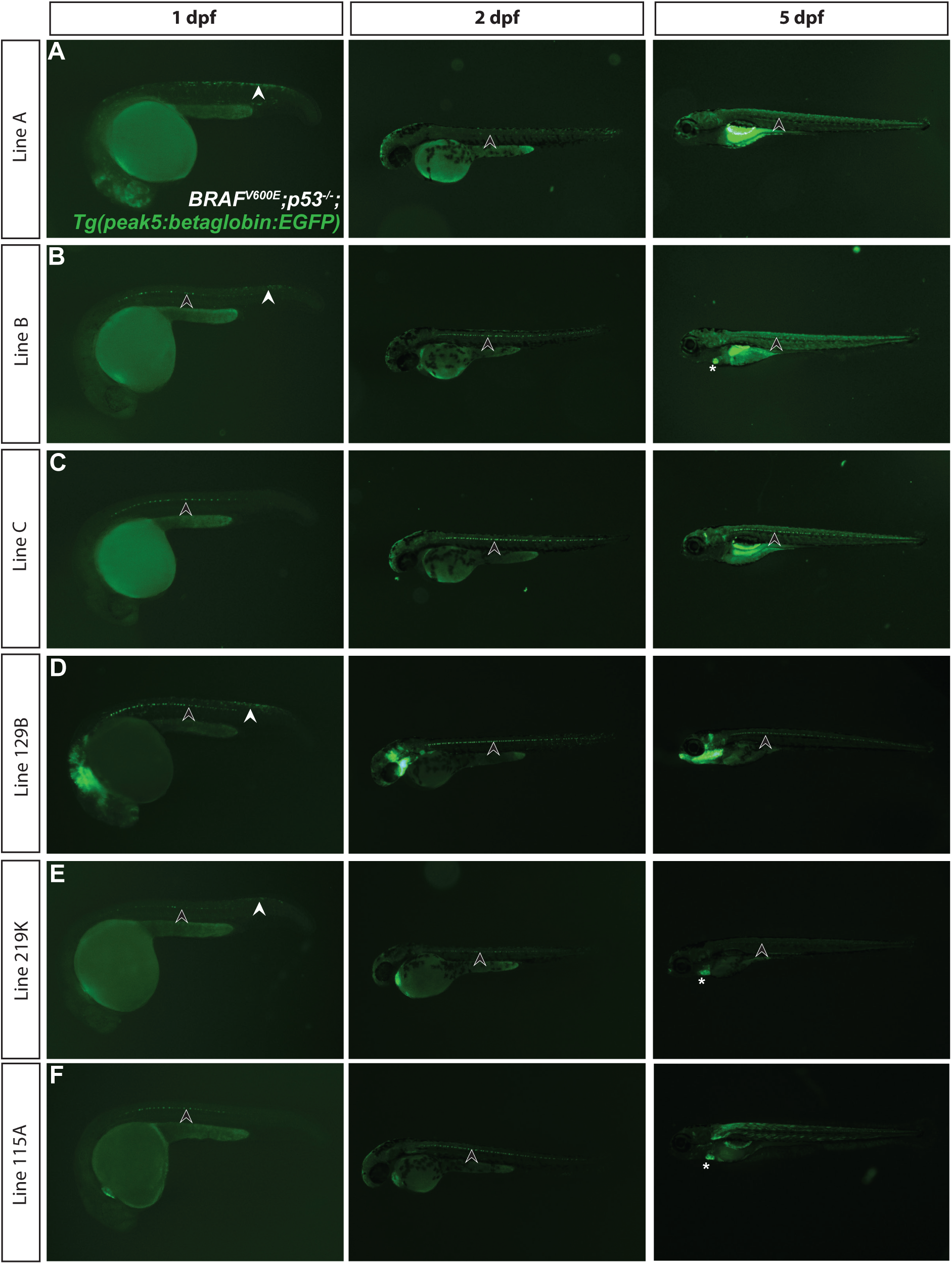
*peak5* stable transgenic lines exhibit similar expression patterns. Multiple transgenic lines derived from independent founder (F0) parental fish were established that transmit the *peak5:betaglobin:EGFP* reporter through the germline. White arrowheads indicate NCCs. Black arrowheads point to KA neurons. Asterisks indicate heart EGFP localization in the heart. **A)** Stable line A exhibits strong EGFP localization in NCCs at 1 dpf. At 2 dpf, some NCCs are labeled, and KA neurons are also faintly visible. At 5 dpf, the most prominently labeled cells are KA neurons and cells near or within the swim bladder. **B)** At 1 dpf, Line B exhibits EGFP localization in posterior premigratory NCCs and strong localization in KA neurons. By 3 dpf and 5 dpf, EGFP localization is primarily within KA neurons and the heart. **C)** Line C mainly exhibits EGFP localization in KA neurons at 1 dpf, 2 dpf, and 3 dpf. **D)** In Line 129B, NCCs, KA neurons, and cells in the cranial CNS are strongly labeled with EGFP at 1 dpf. Labeling of some NCCs is maintained at 2 dpf, and by 5 dpf EGFP is primarily localized within KA neurons and parts of the head. **E)** Line 219K exhibits EGFP localization in some premigratory NCCs, KA neurons, and the heart at 1 dpf. At 5 dpf, EGFP is mainly localized in KA neurons and the heart. **F)** Line 115A displays signs of ectopic *EGFP* expression. At 1 dpf, EGFP localization is mainly visible in KA neurons and the heart. By 2 dpf, EGFP is also localized in the muscles, which is more prominent at 5 dpf. KA neurons and the heart are also labeled at 2 dpf and 5 dpf.

At 1 dpf, 4 out of 6 stable lines exhibited EGFP expression in pre-migratory NCCs dorsally located on the posterior trunk (**Figure 3A, C, D, E**). To confirm that *peak5* is active in NCCs and cells that would typically express *sox10* we crossed *Tg*(*peak5:betaglobin:EGFP*) to either a *Tg*(*crestin:mCh*) line (**Figure 4A’**) or the previously published *Tg*(*sox10(7*.*2):mRFP*) line (Kucenas et al., 2008) (**Figure 4B’**). Epifluoresence imaging of double transgenic embryos at 1 dpf showed most EGFP positive cells in the dorsal posterior region co-localize with mCh or mRFP at 1 dpf (**Figure 4A-B’**), illustrating that *peak5* is active in NCCs. In addition to a subset of NCCs, 6 out 6 *peak5* transgenic lines express *EGFP* within a subset of ventral Kolmer-Agdhur (KA) neurons in the spinal cord, which contact cerebrospinal-fluid (**Figure 3**), as identified by cell shape and localization (**Figure S4**). By 5 dpf, *EGFP* expression was consistently observed in KA neurons across all lines, and in the heart in 3 out 6 stable lines (**Figure 3**).

**Figure 4:**
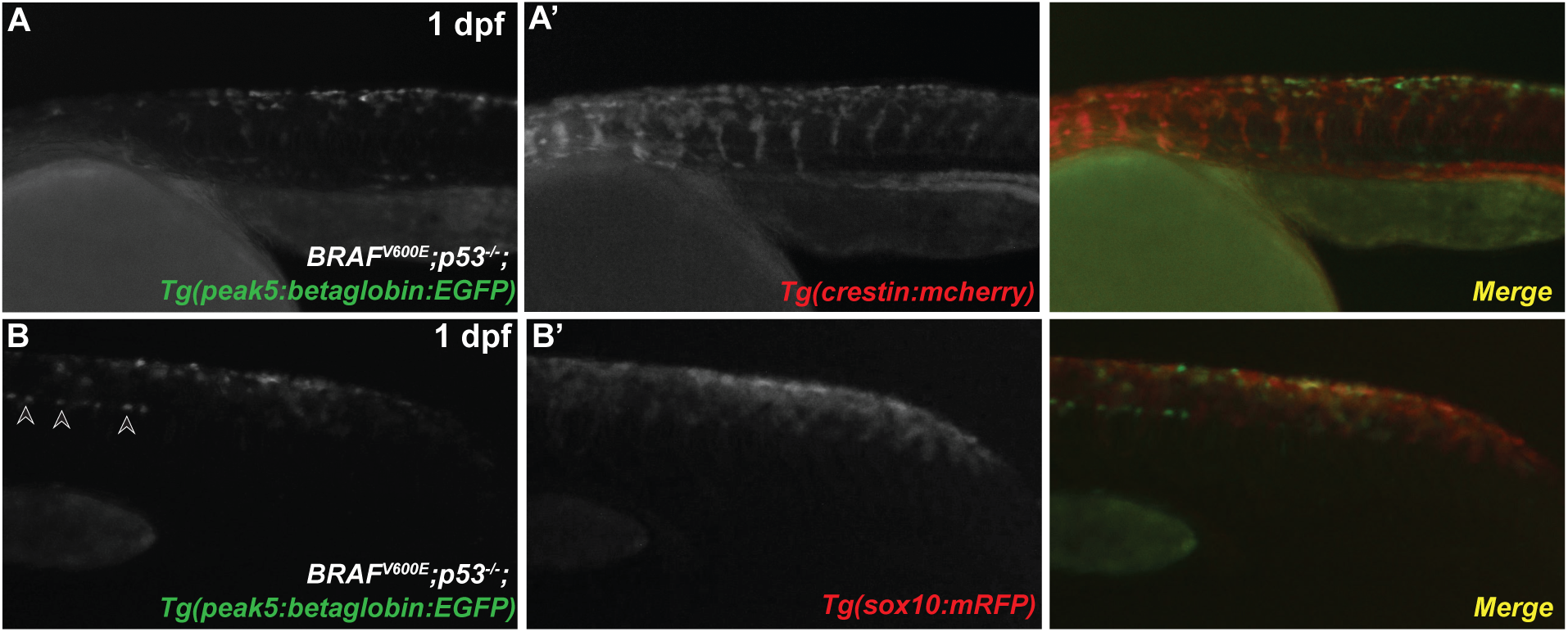
EGFP positive cells co-label with *crestin:mcherry* and *sox10(7*.***2):mRFP***. **A)** *peak5* (Stable Line A) is active in cells with NCC morphology and NCC localization at 1 dpf. **A’)** These EGFP positive cells co-label with *Tg(crestin:mCh)* positive cells. **B)** *peak5* (Stable Line 129B) also exhibits EGFP localization in NCCs and **B’)** co-labels with *Tg(sox10(7*.*2):mRFP)* positive cells. Arrowheads indicate EGFP positive KA neurons.

*EGFP* expression in adult fish was most prominent by epifluorescence in cells in the within the tail, tip of the dorsal fin (**Figure 5A-C**), and peripheral nervous system, particularly observable within the maxillary barbel, a sensory organ, which contains peripheral nerves and neural crest derived Schwann cells (**Figure 5D**). Together, the similar expression patterns of *EGFP* between different stable transgenic lines suggests that our *peak5* transgenic zebrafish reflect endogenous activity of *peak5* in isolation.

**Figure 5:**
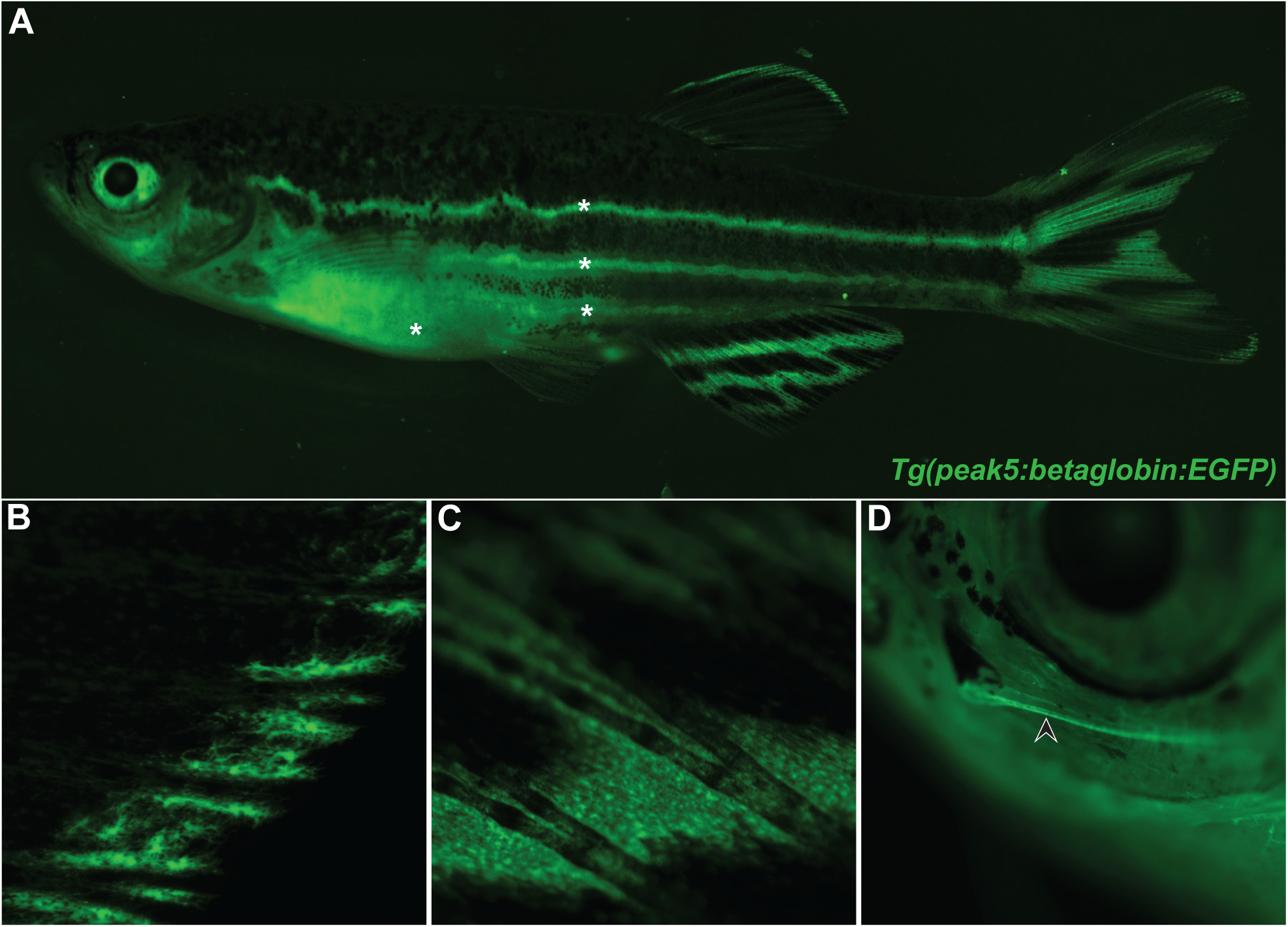
*peak5* is active in regions of fins and the peripheral nervous system in adults. **A)** Whole animal image of *peak5* active in stable transgenic line A. Asterisks (*) indicate regions of auto-fluorescence; a long pass filter set allows discrimination between the yellow hue of autofluorescent tissue and *EGFP*-expressing cells. **B)** EGFP positive cells in the tip of the dorsal fin. **C)** EGFP positive cells in the caudal fin. **D)** EGFP positive cells peripheral nervous system cells within the barbel (arrowhead).

### peak5 is active in early melanoma patches and tumors

Given that *peak5* is active in NCCs, we next asked if *peak5* is an active enhancer in melanoma. Excitingly, in every transgenic line assessed for tumor growth (Lines A, B, and C), *EGFP* was expressed in nearly all melanoma tumors (**Figure 6; Table 4**) (n = 55/59 EGFP+ tumors in 51 animals). Not only is *peak5* active in raised tumors, it is also active in small, unraised, “preclinical” melanoma precursor lesions. Patches of EGFP positive cells, tracked over time, drastically grew (**Figure 7**) (n = 9/15 EGFP+ patches tracked in 13 animals), similar to *crestin:EGFP* positive patches (Kaufman et al., 2016). The remaining 6/15 patches did not obviously expand but remained at least stable over a 2-3 month time period observed. EGFP localization in precursor lesions and tumors is particularly striking because *EGFP* is either minimally or not detectably expressed by epifluorescence in adult melanocytes, relative to patches and tumors, in the lines studied. Of note, differences in expression levels of *EGFP* in the body of the fish may be due to differences in the copy number of the transgene. Overall, *EGFP* expression in patches and tumors indicates that *peak5* is an active enhancer in melanoma, and furthermore, represents a *sox10* regulatory element that is active during early stages of melanoma. To our knowledge, this is the first time a single enhancer has been shown to be active specifically in early melanoma patches and tumors *in vivo*. In addition to *peak5*, we also generated several stable lines to assess activities of the *sox10* minimal promoter, *peak1*, and *peak8* in melanoma. As predicted, the *sox10* minimal promoter drove *EGFP* expression in melanoma patches and tumors across 6 stable lines (n = 38/46 EGFP+ tumors in 42 animals) (**Figure S5A-C’; Table 4**). Unlike *peak5* transgenic lines, *peak1* and *peak8* stable lines expressed *EGFP* embryonically, but did not express *EGFP* in nearly all melanoma tumors (n = 0/6 EGFP+ tumors in 6 *peak1* transgenic animals from 1 stable line; n = 1/26 EGFP+ tumors in 25 *peak8* transgenic animals from 3 independent stable lines) (**Figure S5D-G’; Table 4**). These *peak1* and *peak8* data highlight the specificity of the activity of *peak5* from within the *sox10* super-enhancer and demonstrate that not all sub-elements are sufficient to drive *EGFP* expression in melanoma. These data also show that *peak5* is an enhancer upstream of the *sox10* minimal promoter that is specifically active in melanoma, both in precursor lesions and tumors.

**Figure 6:**
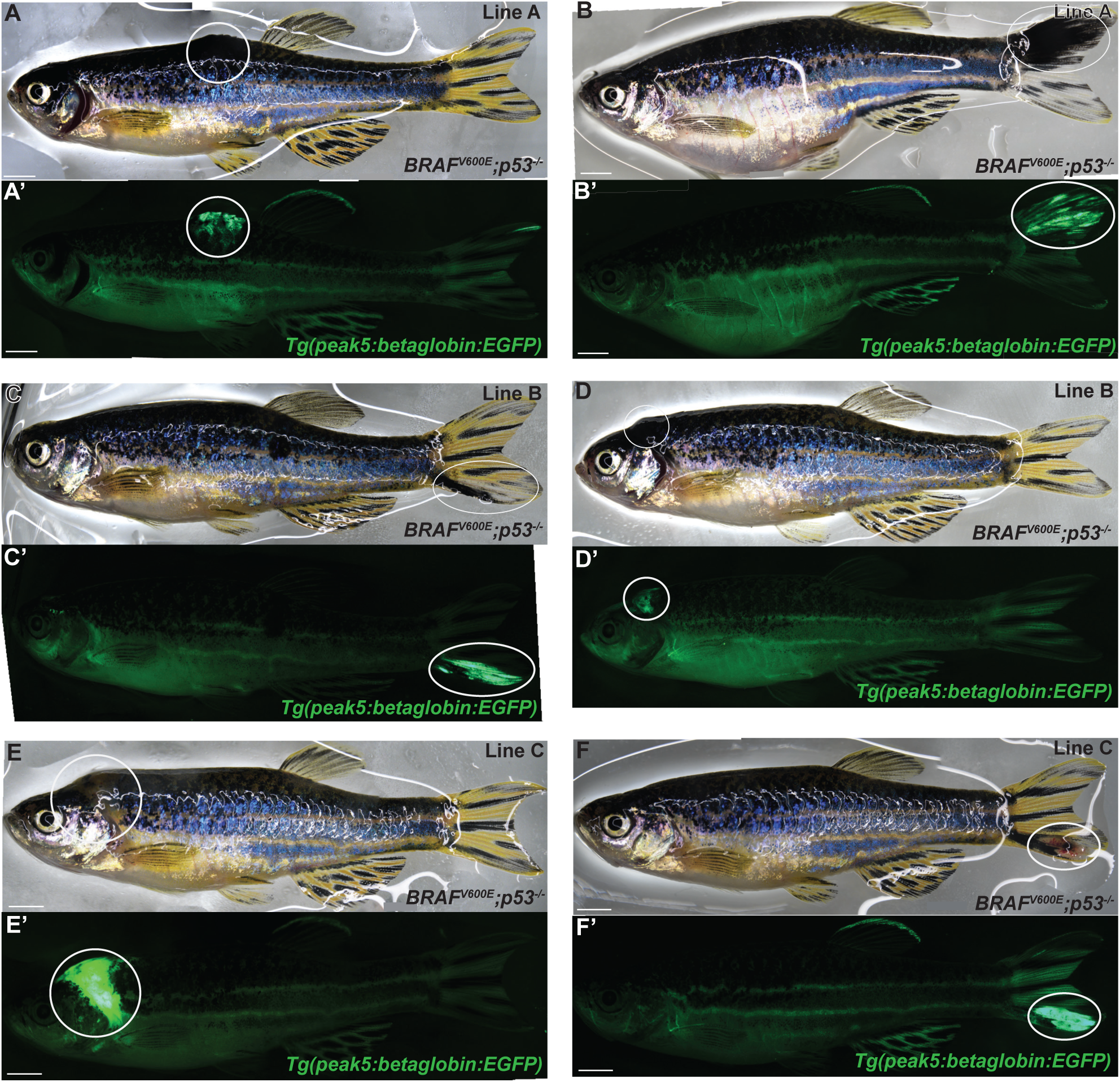
*peak5* is active in melanoma tumors across multiple stable lines. **A-B)** Bright field images of tumors, circled, in *Tg(peak5:betaglobin:EGFP)* stable line A in the *Tg(BRAF*^*V600E*^*);p53*^-/-^ zebrafish background. **A’-B’)** Tumors are EGFP positive and *EGFP* is highly expressed in tumors compared to any low *EGFP* expression elsewhere the fish. **C-D**) Bright field images of *Tg(peak5:betaglobin:EGFP)* stable line B with tumors, circled. **C’-D’)** Tumors are EGFP positive. **E-F**) Bright field images of *Tg(peak5:betaglobin:EGFP)* stable line C with tumors, circled. **E’-F’)** Tumors are EGFP positive.

**Figure 7:**
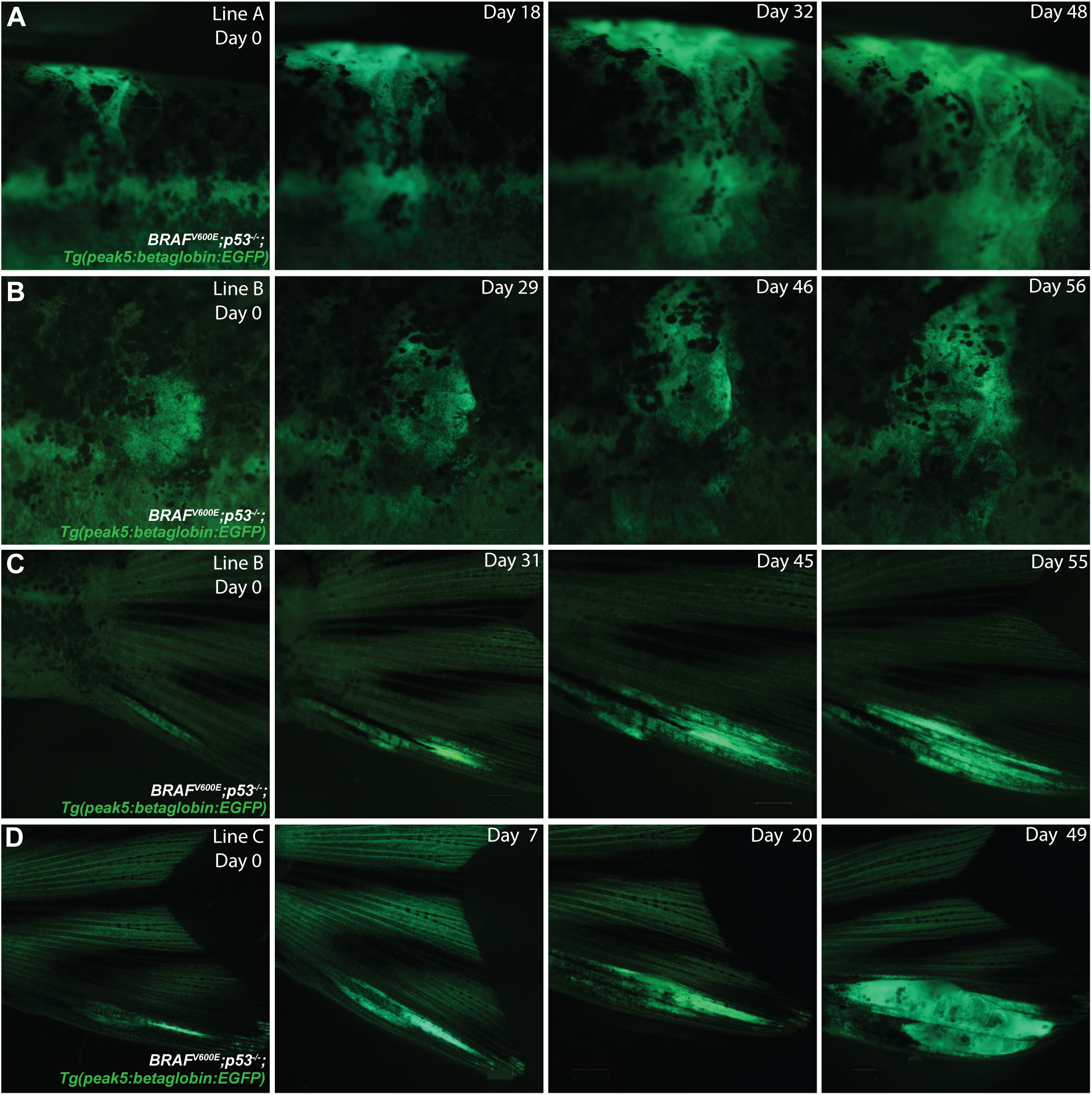
*peak5* is active in melanoma precursor lesions. **A)** *Tg(peak5:betaglobin:EGFP)* stable line A zebrafish with dorsally located melanoma precursor lesion that grows over time. **B)** *Tg(peak5:betaglobin:EGFP)* stable line B zebrafish with a melanoma precursor lesion in a scale that expands in size over time. **C)** *Tg(peak5:betaglobin:EGFP)* stable line B zebrafish with precursor lesion located on the tail that grows over time. **D)** *Tg(peak5:betaglobin:EGFP)* stable line C zebrafish with a precursor lesion on the tail that grows into a tumor. Day 0 denotes the first day the precursor lesion was observed.

### A conserved sequence within peak5 is necessary for activity in NCCs and melanoma

We next asked if within *peak5* there is a minimal sequence that is necessary for activity in neural crest and melanoma. We hypothesized that any nucleotide conservation within *peak5* between zebrafish and other animals would identify a more critical regulatory target region. Nucleotide sequence conservation of enhancers between zebrafish and mammals is rare (Ritter et al., 2010); accordingly, we did not observe any detectable colinear sequence conservation of *peak5* in humans or mice. However, past sequence comparison of the zebrafish exomes to the exomic sequence of its nearest relative, the common carp (*Cyprinus carpio*), revealed high conservation of synteny and homologous coding sequences (Henkel et al., 2012). Thus, we reasoned that evolutionary important non-coding sequences may also be conserved. Aligning the nucleotide sequence around the *sox10* locus in both zebrafish and carp using Nucleotide BLAST revealed discrete regions of conservation, including *peaks 2, 4, 5, 8, 13* and the minimal promoter, as well as most of the *sox10* coding region (**Figure 8A**). Interestingly, these conserved regions correspond to enhancer assays that result in the most robust NCC expression (**Figure S2**).

**Figure 8:**
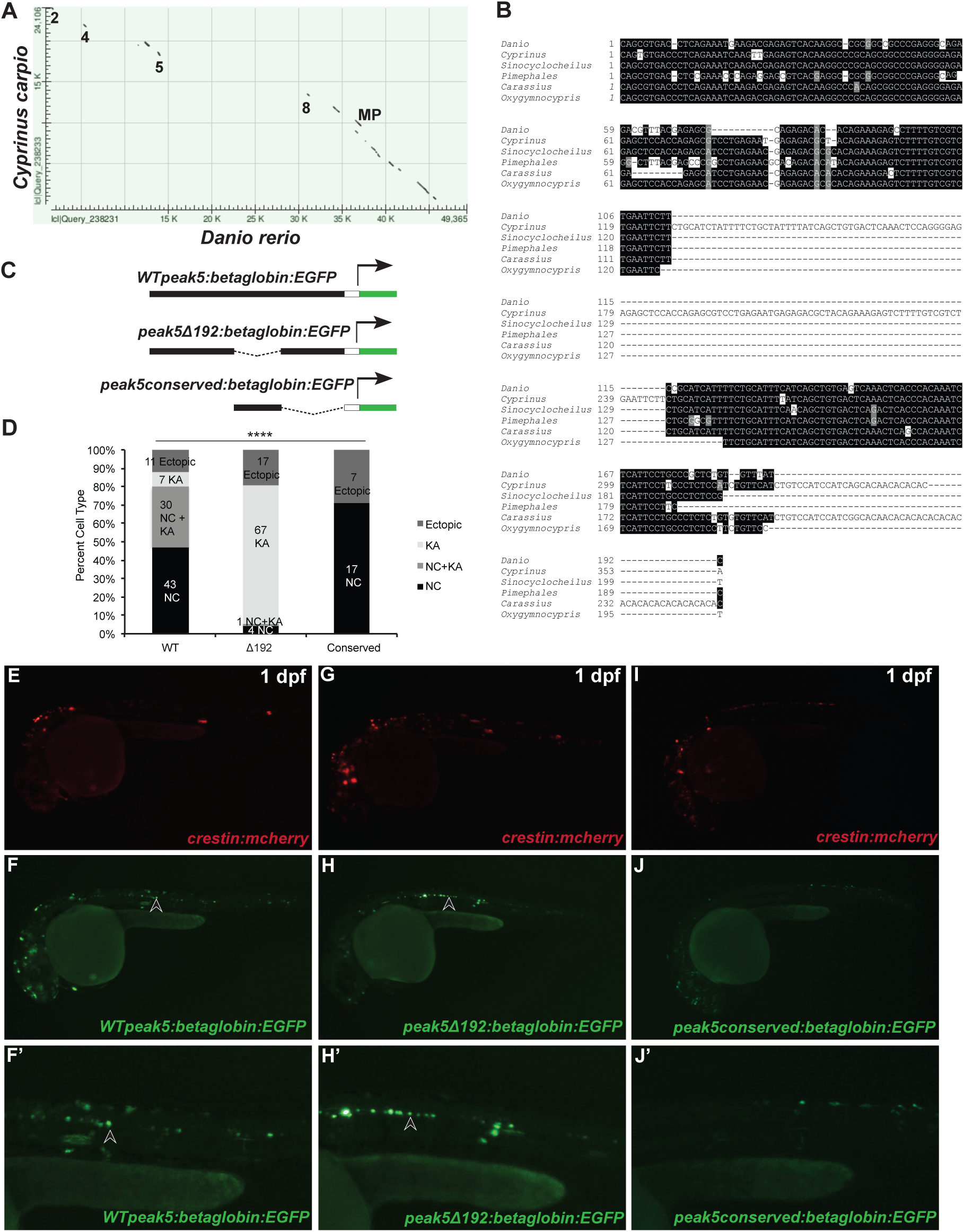
A conserved region of *peak5* is necessary for neural crest activity. **A)** Dot-matrix view of the alignment of the carp (*Cyprinus carpio) sox10* coding and surrounding non-coding genomic region of *sox10* compared to the same genomic region in zebrafish (*Danio rerio*). Numbers indicate regions where peak sequences are conserved. MP = minimal promoter. **B)** Sequence alignment of the most conserved region of *peak5* to members of the *Cyprinidae* family. **C)** Schematic of plasmids used in the experiment. **D)** Quantification of the percentage of categorizable EGFP+ embryos exhibiting NC, KA, or both NC and KA labeling, or ectopic expression. NC = neural crest. KA = Kolmer-Agduhr neurons. **** p-value < 0.0001. **E)** *crestin:mch* expression in an embryo injected with *WTpeak5* and *crestin:mch*. **F)** *WTpeak5* is active in both neural crest and **F’)** KA neurons. **G)** *crestin:mch* expression in an embryo injected with *peak5Δ192* and *crestin:mch*. **H)** *peak5Δ192* is not active in NCCs, but **H’)** is active in KA neurons. **I)** *crestin:mch* expression in an embryo injected with the conserved *peak5* sequence and *crestin:mch*. **J)** The conserved *peak5* sequence is active in neural crest and **J’)** does not exhibit activity in KA neurons. Arrowheads indicate KA neurons.

Within *peak5*, we identified a 192 bp region that is conserved between carp and zebrafish, as well as other selected members of the *Cyprinidae* family (*Sinocyclocheilus grahami, Pimephales promelas, Carassius auratus*, and *Oxygymnocypris stewartii*) (**Figure 8B**). Regions of conservation of *peak5* between species other than carp were located on whole-genome shotgun contigs through Nucleotide BLAST. Therefore, to verify that these regions of conservation are indeed near *sox10* in these other species, we aligned each identified scaffold to the zebrafish *sox10* locus. We identified proximity to the *sox10* locus and/or synteny with other putative *sox10* regulatory regions for all other examined species except *Pimephales promelas* (**Figure S6**).

To test if this *peak5* conserved sequence is required for *peak5* activity, we deleted the sequence within the reporter assay plasmid containing the wild-type *peak5* sequence (**Figure 8C**). Interestingly, compared to full-length wild-type *peak5* sequence, which exhibits activity in NCCs and in KA neurons embryonically, embryos injected with the 192 bp conserved sequence deletion construct (*peak5Δ192:betaglobin:EGFP*) exhibited diminished *peak5* activity in NCCs, while maintaining strong EGFP localization in KA neurons (**Figure 8D-H’**). This suggests that this core conserved sequence controls *peak5* activity within NCCs. Injection of only the *peak5* conserved sequence (*peak5_conserved:betaglobin:EGFP*) resulted in labeling of only NCCs, albeit at weaker levels compared to full-length *peak5*, demonstrating that the conserved sequence is sufficient for *peak5* activity in NCCs and is not sufficient for labeling of KA neurons (**Figure 8D, 8I-J’**).

In accordance with these mosaic analyses, embryos from *peak5Δ192* stable transgenic lines exhibit EGFP localization in KA neurons, but no clear localization in NCCs located in the dorsal posterior region of the trunk (**Figure 9A-A’; Figure S7**). In contrast, NCCs in *peak5_conserved* stable transgenic embryos are EGFP positive and KA neurons are not labeled (**Figure 9B-B’; Figure S7**). In adult stable transgenic *peak5_conserved* zebrafish, cells within the maxillary barbel are labeled, similar to wild-type *peak5* stable lines (**Figure 9D**); however, we did not observe EGFP positive barbels in *peak5Δ192* stable transgenic animals (**Figure 9C**). Furthermore, the majority of tumors in adult *peak5Δ192* stable transgenic animals are not EGFP positive (n = 21/23 EGFP-tumors in 22 animals from 3 different stable transgenic lines) (**Figure 9E-F; Table 4**). These data suggest that the conserved sequence we identified is not only necessary for *peak5* activity in a subset of NCCs, but also is necessary for robust *peak5* activity in melanoma.

**Figure 9:**
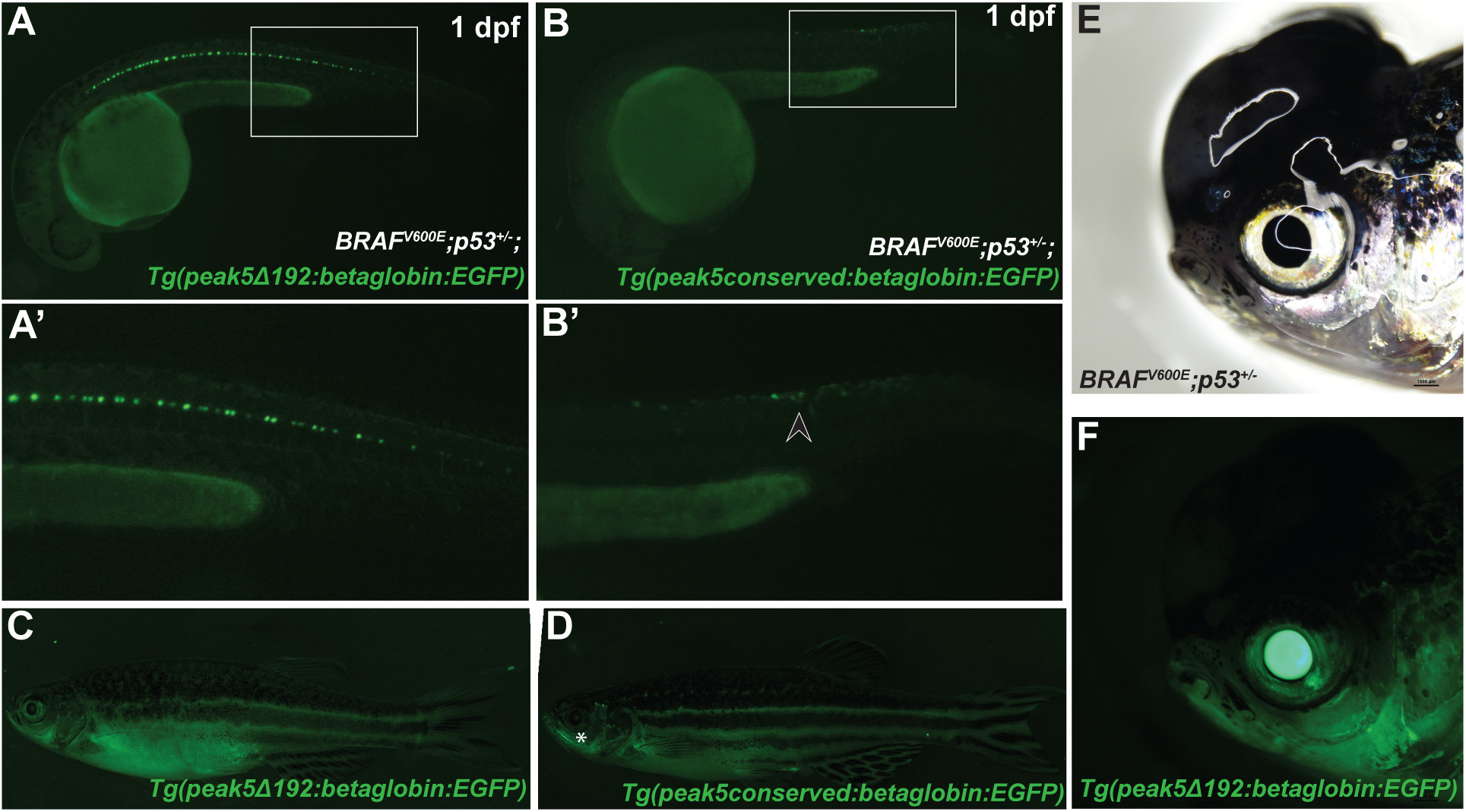
A conserved region of *peak5* is necessary for melanoma activity. **A and A’)** *Tg(peak5Δ192:EGFP)* embryos exhibit EGFP localization in KA neurons, but no overt localization in NCCs in the trunk is observed. **B-B’)** Contrastingly, *Tg(peak_conserved:EGFP)* embryos do not have labeled KA neurons, but exhibit EGFP positive cells in the trunk of the embryo. **C)** Little EGFP localization is observed in *Tg(peak5Δ192:EGFP)* adult fish whereas **D)** cells within the maxillary barbels (asterisk) of *Tg(peak_conserved:EGFP)* are EGFP positive. **E-F)** The majority of tumors that develop in *Tg(peak5Δ192:EGFP)* are EGFP negative.

### SoxE TFBS regulate peak5 activity in NCCs

We next asked which TFBS within this conserved region of *peak5* are functionally necessary for activity. Out of a list of predicted TFBS, some of which have roles in NCC development, our attention was drawn to predicted dimeric SoxE binding sites (**Figure 10A**) as it was previously noted that such dimeric SoxE binding sites have been shown to exhibit functional relevance in mouse *Sox10* enhancers (Antonellis et al., 2008). Furthermore, dimeric SOXE TFBS with 3-5 bp spacers are enriched in human melanoma cell putative enhancers identified by DNase hypersensitivity site mapping (Huang et al., 2015). We therefore mutated both of these predicted SoxE binding sites within the *peak5* WT sequence to functionally test their necessity (**Figure 10B-C**). Mutated plasmid was co-injected with *crestin:mch* and scored for activity in NCCs and KA neurons. Compared to *peak5* wild-type injected embryos, mutation of the SoxE binding sites result in drastically diminished *EGFP* expression in NCCs, but KA *EGFP* expression is maintained (**Figure 10D-J**). This mutational analysis confirms that the dimeric SoxE binding sites are functional and likely important for the transcriptional regulatory activity of *peak5* in the neural crest.

**Figure 10:**
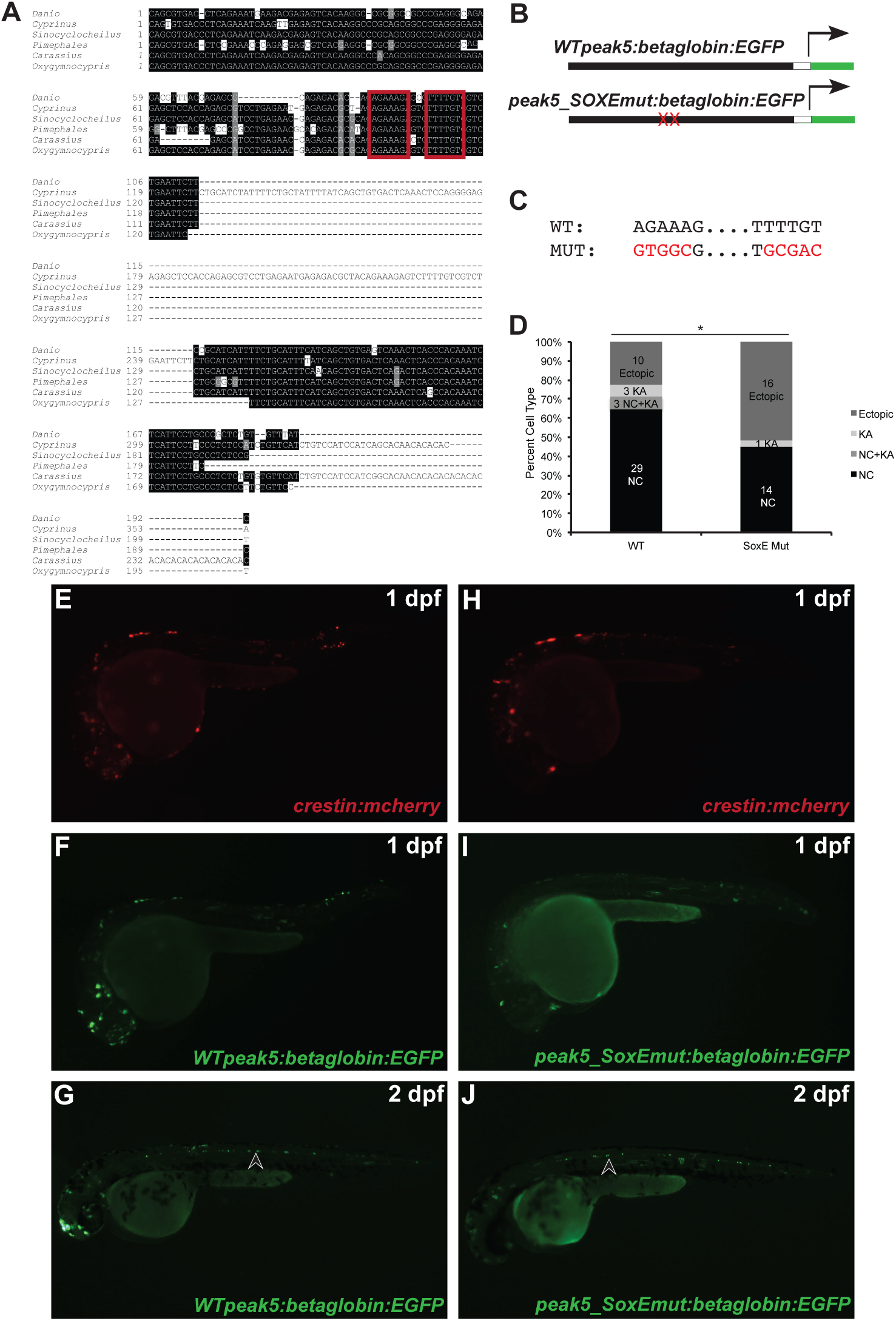
Mutation of dimeric SoxE TFBS affect *peak5* neural crest activity. **A)** Dimeric SoxE TFBS, as predicted by JASPAR (outlined in red), are present in the conserved sequence of *peak5*. **B)** Schematic of plasmids used in the experiment. Mutations to SoxE TFBS were made in the context of the full-length WT *peak5* plasmid. **C)** Five out of six nucleotides for each motif were mutated. **D)** Quantification of the percentage of categorizable EGFP+ embryos exhibiting NC, KA, both NC and KA, or ectopic expression. NC = neural crest. KA = Kolmer-Agduhr neurons. * p-value = 0.0398. **E)** *crestin:mch* expression in an embryo injected with *WT peak 5* and *crestin:mch*. **F)** *WT peak5* is strongly active in neural crest at 1 dpf. **G)** *WT peak5* is strongly active in neural crest derived cells and KA neurons at 2 dpf. **H)** *peak5_SoxEmut* is not active in the neural crest, but **J)** is active in KA neurons. Arrowheads indicate KA neurons.

## Discussion

Melanoma initiating cells express aspects of a neural crest program that is maintained as a tumor progresses (Kaufman et al., 2016). One gene within the neural crest program, *sox10*, is a critical regulator of NCC development and is also upregulated in both zebrafish and human melanomas. Indeed, human melanomas immunostain positive for Sox10 in most cases, even more often than the key melanocyte lineage regulator MITF (Clevenger et al., 2014; Ordóñez, 2014; Willis et al., 2015). Past work demonstrated that modulation of *sox10* expression in melanocytes, which normally lowly express *sox10*, affects melanoma onset rates (Clevenger et al., 2014; Kaufman et al., 2016; Shakhova et al., 2012), illustrating that *sox10* plays a key role during melanoma initiation. Therefore, deciphering the epigenetic regulation of *sox10* may elucidate earlier mechanisms of melanoma initiation that drive NCP activation. Ongoing studies continue to decipher the embryonic function of *sox10* in zebrafish (Chong-Morrison et al., 2018; Dutton et al., 2001; Kelsh and Eisen, 2000), but to date, regulation of *sox10* expression in melanoma is poorly understood. In this study, we screened individual putative *sox10* regulatory elements for activity within NCCs and melanoma in zebrafish. From this analysis, we identified *peak5*, an enhancer that is active in embryonic neural crest and melanoma, including precursor lesions.

Using ATAC-Seq data generated from zebrafish melanoma tumors, we identified putative regulatory regions of open chromatin surrounding and within the *sox10* locus. Injection of reporter constructs harboring these putative enhancers into embryos revealed that 9 out of 11 displayed some level of activity within NCCs, as compared to injected negative controls, indicating their potential as active enhancers in melanoma. Generation of multiple stable lines revealed that *peak5*, a 669 bp sequence from 22.8 kb upstream of the *sox10* transcriptional start site, is active embryonically in a subset of NCCs and KA neurons. In zebrafish adults, *peak5* is at most minimally active in melanocytes in *in vivo* EGFP reporter assays, and is highly active in melanoma patches and tumors. Furthermore, we identified a region within *peak5* that is conserved between members of the *Cyprinidae* family and mediates the NCC activity of *peak5*. Within this conserved region, dimeric SoxE binding sites are critically necessary for *peak5* activity within the neural crest. Together, *peak5* serves as a proxy enhancer to understand how transcriptional regulation of *sox10* and potentially other melanoma enriched genes influences melanoma initiation.

The identification of *crestin* as an early marker of melanoma provided a live marker of the earliest detectable events of melanoma initiation and confirmed that the NCP activation is an early step in the process (Kaufman et al., 2016). This study expands upon this knowledge by identifying a regulatory element that can also be utilized as an *in vivo* specific reporter of melanoma initiation. Importantly, unlike *crestin, sox10* is conserved across all vertebrate species; therefore, identifying *sox10* enhancers that are active in melanoma in zebrafish could lead to the identification of human *SOX10* enhancers that influence melanoma onset. While we were unable to identify stretches of colinear conserved sequence between the zebrafish and human/mouse upstream regions of *SOX10*, predicting the regulatory function of enhancers based on sequence conservation alone is often not possible, such as in the case of *RET* enhancer elements, which function faithfully between human and zebrafish in reporter assays, but are not conserved in an obvious way at the primary nucleotide level (Fisher, 2006).

Further, we were intrigued to find that while zebrafish *sox10* enhancers are not conserved in a colinear sequence with potential human *SOX10* enhancers, several zebrafish enhancers are conserved by nucleotide sequence between members of the *Cyprinidae* family. Conservation of *peaks 2, 4, 5, 8, 13* and the *sox10* minimal promoter between zebrafish and carp suggest an evolutionary pressure to maintain these regulatory sequences. Interestingly, the highest region of conservation for each peak is oriented near the center of the ATAC-Seq peak, including *peak5*. As the conserved region of *peak5* controls activity in NCCs, but not activity in KA neurons, regions of conservation in *peaks 2, 4, 8*, and *13* may represent core sequences that control NCC activity of the enhancer. Future work to dissect the role of these core conserved enhancers in NCC development and melanoma will test this hypothesis.

This region of high conservation within *peak5* also leads to the hypothesis that the key TFBS controlling *peak5* activity in NCCs and melanoma are also conserved. Identification of SoxE dimeric binding sites points toward potential transcription factors, most notably Sox10, that may bind and activate *peak5*. Similar to deletion of the conserved sequence within the context of full-length *peak5*, mutation of both SoxE TFBS lessens *peak5* activity in NCC, but does not affect *peak5* activity within KA neurons. Therefore, members of the SoxE family (Sox10, Sox9, and Sox8) are candidate transcription factors that may bind and activate *peak5*. Dimeric SOXE binding sites are overrepresented in human melanoma cells in regions of open chromatin (Huang et al., 2015) and have also been observed near promoters of genes regulated by Sox10 (Mollaaghababa and Pavan, 2003), suggesting that Sox10 may bind the dimeric sites in *peak5*. Mutation of the dimerization domain within Sox10 in mice also causes defects in melanocyte development, underscoring the importance of dimeric Sox10 binding (Schreiner et al., 2007).

Interestingly, dimeric SoxE sites have also been shown to be necessary for activity of two mouse *Sox10* enhancers, MCS4 and MCS7 (Antonellis et al., 2008). Since zebrafish *sox10* enhancers are not conserved by sequence with mouse *Sox10* enhancers, it will be interesting to test if either mouse MCS4 or MCS7 *Sox10* enhancers have overlapping embryonic and adult activity patterns with *peak5* in zebrafish. Moreover, examining the other functional TFBS present in *peak5* in zebrafish may help predict and identify a *peak5*-equvialent enhancer in humans that is depend on the same or similar complement of TFBS.

One caveat of determining the functionality of a putative enhancer is the potential to identify false positives. In zebrafish, random genome integration of enhancers using the Tol2 system can result in enhancer trapping (Udvadia and Linney, 2003). Thus, we generated multiple stable lines for *peak5*, which exhibited similar expression patterns, suggesting that our lines reflect endogenous spatial and temporal activity of *peak5* embryonically and in melanoma. Variations amongst transgene integration site in different stable lines leading to subtle position effects in addition to genomic instability in melanoma tumors, may also explain why we occasionally observe EGFP negative tumors in stable lines that predominately express EGFP in tumors (*peak5* n = 4/59 EGFP negative tumors) and a small number of EGFP positive late-stage tumors in *peak8* (n = 1/26) and *peak5Δ192* (n = 2/23) stable lines.

Finally, we propose a model by which *peak5* contributes to increased *sox10* expression regulation in melanoma initiation and progression (**Figure 11**). In this model, the *peak5* region is open and enhances *sox10* expression in early melanoma precursor lesions and, through a feed-forward mechanism, autoregulates increasing *sox10* expression by binding at the dimeric SoxE sites in *peak5*. Again, while stretches of conserved, noncoding DNA are most often not readily identifiable by comparison between humans/mouse and zebrafish, the presence of such dimeric SoxE sites in *bona fide* murine NCC enhancers (Antonellis et al., 2008) and putative human melanoma enhancers (Huang et al., 2015) is consistent with this potentially being a general mechanisms by which Sox10 target genes are upregulated in melanoma. Given that *peak5* is not active in all cells that express *sox10* during embryogenesis, other *sox10* enhancers likely also contribute to *sox10* regulation during melanoma initiation. Together, developmentally incorrect re-activation of *sox10* enhancers in zebrafish and human melanomas suggests that understanding the activity and contribution of individual enhancers within this locus will illuminate earlier initiating steps of melanoma. This study identifies one of these enhancers, *peak5*, that is active embryonically and in melanoma, and lays the groundwork to further dissect the role of other *sox10* enhancers in zebrafish and higher vertebrates in melanoma initiation.

**Figure 11:**
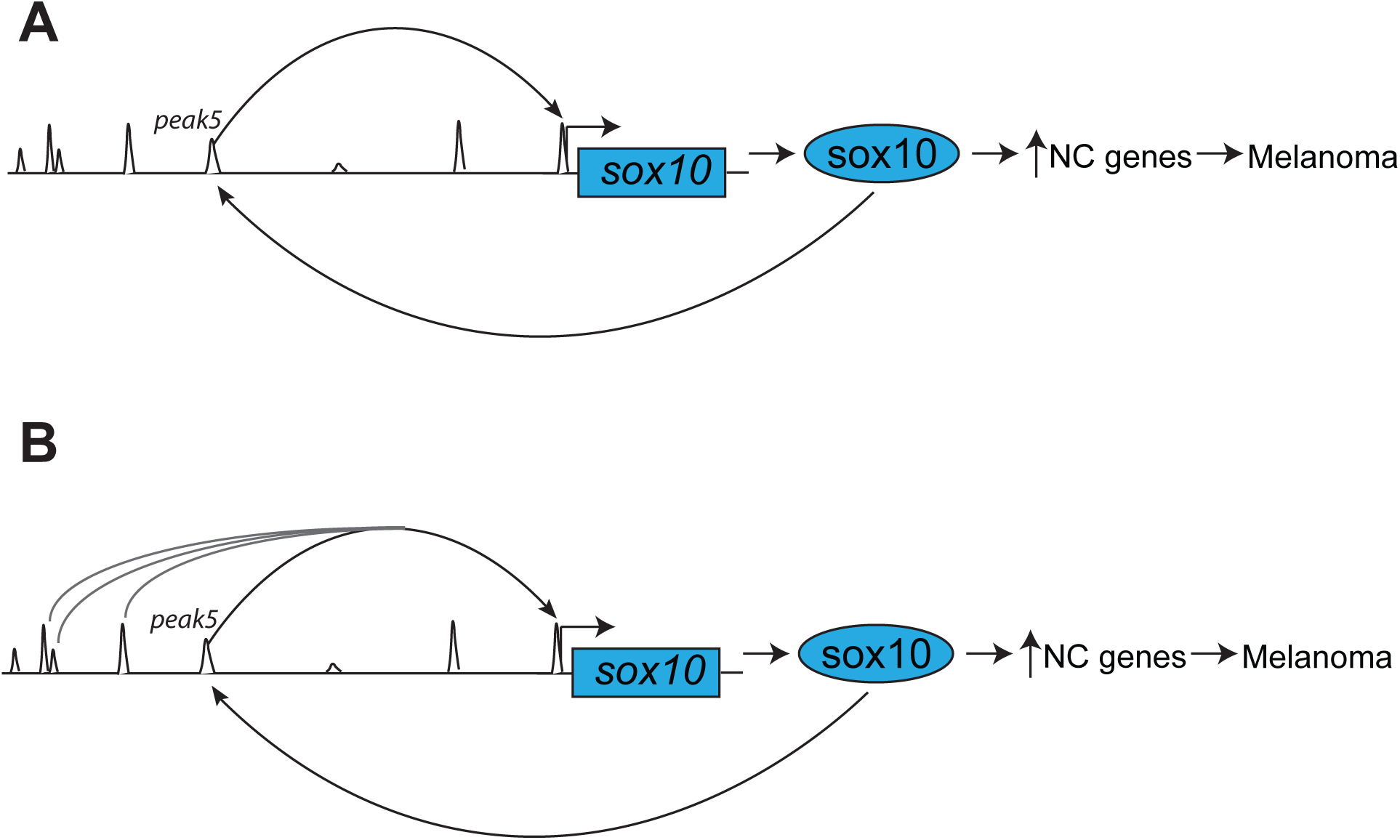
Model of *peak5* regulation of *sox10* expression during melanoma initiation. **A)** In our model, *peak5* activates *sox10* expression in NCCs during development and then again in melanoma precursor cells, likely through dimeric SoxE TFBS. This leads to upregulation of neural crest related genes and melanoma onset. **B)** *peak5* is most likely not the only *sox10* enhancer that regulates *sox10* expression. Multiple elements within the *sox10* super-enhancer may coordinate together to upregulate *sox10* expression during melanoma initiation.

## Methods and Materials

### Zebrafish Lines and Rearing Conditions

Zebrafish were reared in accordance with Washington University IACUC animal protocols in the Washington University Zebrafish Consortium Facility. Adult zebrafish were crossed either as pairs or groups, and embryos were raised in egg water (5 mM NaCl, 0.17 mM KCl, 0.33 mM CaCl_2_, 0.33 mM MgSO_4_) at 28.5°C. Larvae were staged at days post fertilization (dpf) and adults were staged at months post fertilization (mpf). The following strains of transgenic zebrafish were used in this study: AB*, *Tg(BRAF*^*V600E*^*/p53*^*-/-*^*/crestin:EGFP)* (Kaufman et al., 2016), *Tg(BRAF*^*V600E*^*);p53*^*-/-*^ (Patton et al., 2005), *Tg(sox10(7*.*2):mRFP)* (Kucenas et al., 2008), *Tg(crestin:mcherry), Tg*(*peak5:betaglobin:EGFP*), *Tg*(*peak5Δ192:betaglobin:EGFP*), *Tg*(*peak5_conserved:betaglobin:EGFP*), *Tg*(*sox10minimalpromoter:EGFP*), *Tg(peak1:betaglobin:EGFP*), and *Tg*(*peak8:betaglobin:EGFP*).

### Isolation of Bulk Tumors

Following humane euthanasia, grossly visible bulk melanoma tumors from *Tg(BRAF*^*V600E*^*/ crestin:EGFP);p53*^*-/-*^ zebrafish were excised with a razor, manually sheared, and incubated in fresh 0.9X PBS with 2.5 mg/mL Liberase for up to 30 minutes to dissociate cells. Fetal bovine serum addition terminated the digestion, and the cells were passed through a 40 mm filter. Cells were centrifuged at 2000 × g for 5 minutes at 4°C. Supernatant was removed and the cells were resuspended in 500 µL of 0.9X PBS and kept on ice.

### ATAC-Seq

50,000 cells per sample underwent tagmentation reaction with Nextera Tn5 transposase using the Illumina Nextera kit and purified with a Qiagen MinElute reaction kit (Buenrostro et al., 2015). The DNA was then PCR amplified for 9 cycles to add indexing primers. SPRI AMPure beads enriched for fragments under ∼600 bp. The DNA was cycled again with the 9-cycle protocol, followed by cleanup with SPRI AMPure beads. The DNA was quantified with a Qubit DNA High Sensitivity assay and analyzed for quality and size distribution on an Agilent TapeStation with a High Sensitivity D5000 ScreenTape. Samples were pooled for a 10 nM final overall concentration. Sequencing was performed with an Illumina HiSeq 2500 system with 2 × 50 bp read length by the Washington University in St. Louis School of Medicine Genome Technology Access Center.

### ATAC-Seq Data Analysis

Reads were demultiplexed then trimmed with Cutadapt (adaptor sequence 5’-CTGTCTCTTATACACATCT-3’ for both reads) and checked with FastQC to ensure quality. The reads were aligned to the GRCz10/danRer10 genome using BWA-MEM and sorted using SAMtools. Duplicate reads removed with Picard tools using the following parameters: ASSUME_SORTED=true, VALIDATION_STRINGENCY=LENIENT. The files were indexed with SAMtools then filtered for high quality alignments using the following parameters: -f 3, -F 4, -F 8, -F 256, -F 1024, -F 2048, -q 30. MACS2 was then used to identify peaks with the callpeak command with parameters -g 1.4e9, -q 0.05, --nomodel, --shift -100, --extsize 200. Differential peak accessibility was assessed with the MACS2 bdgdiff function. HOMER was then used to annotate the peaks (annotatePeaks.pl) and identify enriched transcription factor motifs using findMotifsGenome.pl (-size 200, -mask parameters).

### Cloning putative enhancers

All putative enhancer elements except for *peak13* were PCR amplified from genomic DNA from AB* zebrafish larvae or *Tg*(*BRAF*^*V600E*^)*;p53*^*-/-*^ larvae with high-fidelity DNA polymerases. *peak13* was amplified from BAC CHORI-211:212I14 (a gift from the Johnson Lab). Genomic locations for each putative enhancer in the Zv10 build of the zebrafish genome are listed in **Table 1** and primers used to amplify each sequence and total cloned sizes of each element are listed in **Table 2**. PCR amplified enhancer elements were cloned into pENTR5’ vectors (ThermoFisher). Anytime Phusion (NEB) was utilized to amplify a putative enhancer, adenines were added to the end of the PCR product using *Taq* polymerase (Promega) before performing a TOPO reaction. Using plasmids from the Tol2 kit, all putative enhancers were placed upstream of a basal promoter, mouse *beta-globin* (Tamplin et al., 2011) via Gateway LR reactions with the following combination of Gateway vectors: p5E-enhancer, pME-*betaglobin:EGFP*, p3E-polyA, and pDestTol2pA. To generate a *sox10* minimal promoter plasmid, a *sox10* minimal promoter was PCR amplified from genomic DNA from AB* larvae with primers (**Table 1 and 2**) containing overlapping nucleotides to sequences on either side of the AgeI site in pENTR-EGFP2 (Addgene #22450), similarly described in (Quillien et al., 2017). The PCR product was then cloned into pENTR-EGFP2 and digested with AgeI, through a Gibson Assembly reaction using an in-house Gibson mixture (gifted by the Solnica-Krezel lab, Washington University in St. Louis; NEB Phusion, NEB Taq DNA Ligase, NEB T5 Endonuclease). This pME-*sox10minimalpromoter:EGFP* was then used in a Gateway LR reaction with p5E-MCS, p3E-polyA, and pTol2Destp2A to generate the final reporter construct. All PCR amplified sequences were Sanger sequenced by Genewiz. The list of plasmids used in this study can be found in **Table 3**. The conserved *peak5* sequence was PCR amplified from the previously cloned *peak5* sequence, cloned into a pENTR5’ vector, and placed upstream of *betaglobin:EGFP* through a Gateway reaction, as described above.

### Screening putative enhancers and generation of transgenic zebrafish lines

One cell stage *Tg(BRAF*^*V600E*^*);p53*^*-/-*^ embryos were injected with 20 ng/µL of an enhancer reporter plasmid in addition to 15 ng/µL of *crestin:mch*, and 20 ng/µL of Tol2 transposase mRNA. 1 dpf larvae were screened for EGFP and mCh localization in the neural crest on a stereomicroscope. Embryos were scored as “strong expression” if 5 or more cells were EGFP and mCh positive. Three technical and biological replicates of injections were performed for each putative enhancer. At least 30 embryos scored as positive from each replicate were raised to adulthood. Founders were identified by out-crossing F0 adults, and progeny were screened from 1 dpf - 5 dpf for EGFP localization and subsequently raised to adulthood and screened for *EGFP* expression in melanoma patches and tumors (**Table 4**).

### Imaging

For live-imaging, larvae were anesthetized in Tricaine, embedded in 0.8% agarose, and imaged with a Nikon SMZ18 fluorescent dissecting microscope. For imaging adults, fish were anesthetized in Tricaine and imaged with a Nikon SMZ18 fluorescent dissecting microscope. Images were processed with Photoshop and ImageJ. The Photomerge function in Photoshop was used to stitch together tiled images of adult zebrafish.

### Evolutionary conserved sequence identification

To generate the Dot-Matrix view, sequences surrounding the *sox10* locus in both zebrafish (Chromosome 3: 1976387-2026723, danRer10) and carp (scaffold: LG6, Chromosome 6: 16126287-16150639), were aligned with optimization for more dissimilar sequences in BLAST (http://blast.ncbi.nlm.nih.gov). BLAST with optimization for more dissimilar sequences with discontiguous megablast was also used to compare the zebrafish *peak5* sequence to selected members of the *Cyprinidae* family (**Table 5**). The nucleotide collection or whole-genome shotgun contigs databases were chosen as the search set. The most conserved regions of *peak5* nucleotide sequences from each species were then aligned with M-Coffee (http://tcoffee.crg.cat/apps/tcoffee/do:mcoffee). The fasta_aln file was downloaded and Boxshade (https://embnet.vital-it.ch/software/BOX_form.html) was then used to shade the alignment using RTF_new as the output format and “other” as the input sequence format. TFBS predictions were identified with FIMO (http://meme-suite.org/tools/fimo) using the JASPAR core 2016 vertebrate list. SoxE dimeric binding sites were predicted with JASPAR (http://jaspar.genereg.net/search?q=&collection=CORE&tax_group=vertebrates), searching explicitly for Sox10 motifs using an 80% relative profile score threshold.

### Plasmid Mutagenesis and Deletion Analysis

The Q5 Site-Directed Mutagenesis Kit (NEB) was used according to the manufacturer’s instructions to either delete the *peak5* conserved 192 base pair sequence or mutate SoxE binding sites. The correct deletion or mutations were confirmed through Sanger Sequencing at Genewiz. 20 ng/µL of *peak5* full-length wild type, *peak5Δ192*, or *peak5* full-length SoxE site mutated plasmids were co-injected with 15 ng/µL of *crestin:mch* and 20 ng/µL of Tol2 transposase mRNA into embryos derived from a *Tg(BRAF*^*V600E*^*);p53*^*-/-*^ in-cross. Only embryos that expressed *mCh*, ensuring successful injection, were scored on 1 dpf for expression of *EGFP*. EGFP+ embryos that had identifiable cell type labeling were categorized into the following cell-localization categories: neural crest, KA neurons, both the neural crest and KA neurons, or ectopic. The scorer was blinded to treatment groups when scoring. Three technical and biological replicates were performed for the conserved deletion experiment and two technical and biological replicates were performed for the SoxE TFBS mutation experiment. Stable lines were identified and characterized as described above (**Table 4**).

### Statistical Analyses

Chi-squared analyses were used to compare *peak5* conserved sequence deletion and SoxE TFBS mutagenesis injection experiments in GraphPad Prism 8.

## Author Contributions

R.L.C., E.T.K, and C.K.K. designed research, and R.L.C, E.T.K., S.K.D, V.G., and S.S. performed research. R.L.C. and C.K.K. analyzed data, and R.L.C. and C.K.K. wrote the manuscript.

## Acknowledgements

Research reported in this publication was supported in part by the National Cancer Institute of the National Institutes of Health under award number R01CA240633. The content is solely the responsibility of the authors and does not necessarily represent the official views of the National Institutes of Health. R.L.C. was supported by the National Science Foundation Graduate Research Fellowship (DGE-1745038). C.K.K. was supported by the Cancer Research Foundation Young Investigator Award. E.T.K was supported by T32 GM007067, and S.K.D is supported by T32 GM007067. We thank members of the Kaufman lab and Souroullas lab for helpful discussions; the Washington University Zebrafish Consortium; the Washington University in St. Louis School of Medicine Genome Technology Access Center; Bo Zhang in the Department of Developmental Biology and the Center for Regenerative Medicine at Washington University School of Medicine for assistance with ATAC-Seq analysis; S. Johnson lab members for the BAC plasmid (Washington University in St. Louis); S. Kucenas (University of Virginia) for the *Tg(sox10:mRFP)* line; M. Bagnall (Washington University in St. Louis) for identifying the Kolmer-Agduhr neurons.

