## Supplemental Figures and Tables for "Functional *in vivo* characterization of zebrafish *sox10* enhancers in melanoma and neural crest"

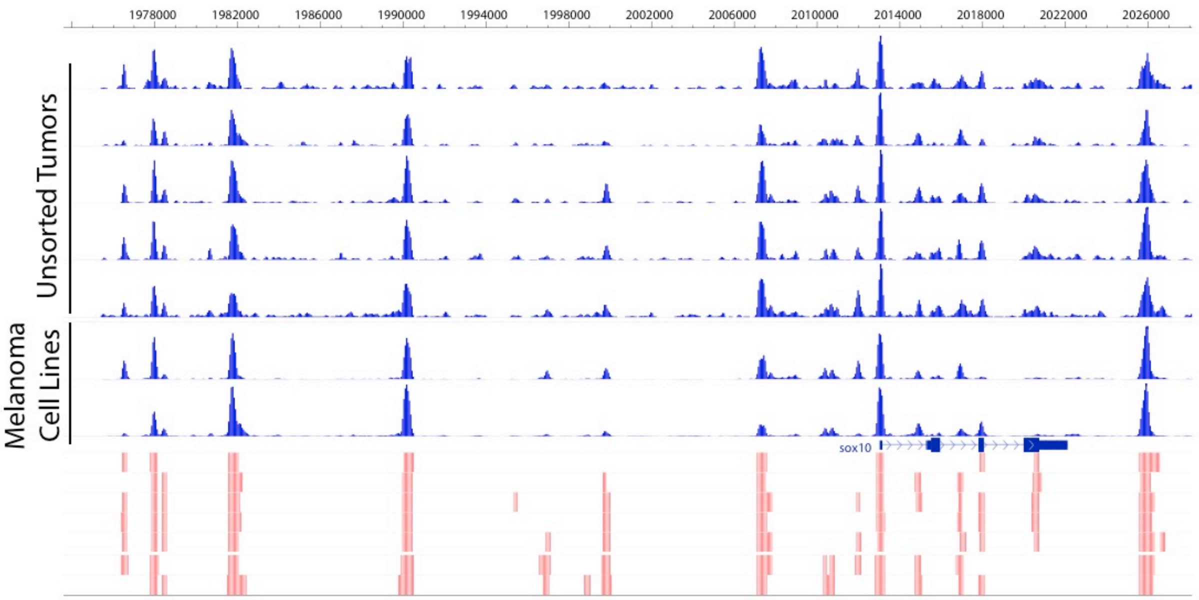


**Figure S1: ATAC-Seq peaks are consistent across tumors**

ATAC-Seq peaks surrounding the zebrafish *sox10* locus. Zebrafish melanoma cell line ATAC-Seq data (Kaufman et al., 2016) aligns with tumor sample data. Red bars indicate potential peaks.

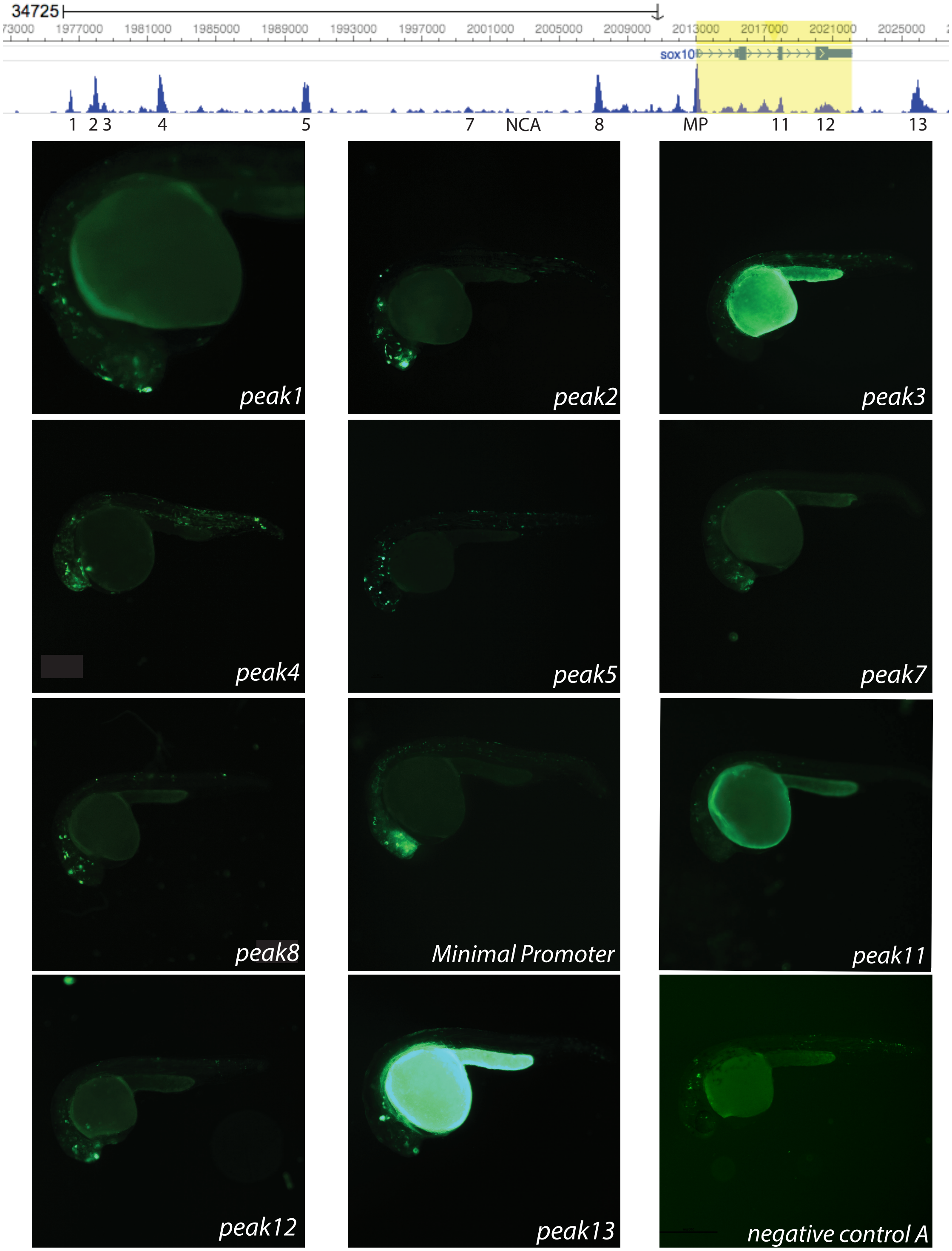


**Figure S2: 9 out of 11 peaks are active embryonically**

**A)** Regions of open chromatin within the *sox10* coding and non-coding genomic locus. NCA = negative control, MP = minimal promoter. **B)** 1 dpf images of F0 mosaic embryos expressing putative *sox10* enhancer assay reporters.

```
Enhancer5_sox10.ape from 735 to 1529
Alignment to
peak5_cloned.ape--  Matches:661; Mismatches:8; Gaps:126; Unattempted:0

             *         *         *         *         *         *         *         *         *         *
    735>agtcaatctgacagggtattgttgagtttttggaaatattcaaagcattttcccaaaatatgtgtcaaaacaagatttgtcaccaaaaatcatcctattt>834
      1>agtcaatctgacagggtattgttgagtttttggaaatattcaaagcattttcccaaaatatgtgtcaaaacaagatttgtcaccaa aaatcatcctattt>100

             *         *         *         *         *         *         *         *         *         *
    835>gtggtcaaaataactcttaaaattccccagagctgatgaaaacgacccaacagcacacacacgctacagtgttttcctctggacagactgactgggtgat>934
    101>gtggtcaaaataactcttaaaattccccagagctgatgaaaacgacccaacagcacacacacgctacagtgttttcctctggacagactgactgggcgat>200

             *         *         *         *         *         *         *         *         *         *
    935>GTGAAAGCTCTCTGCACCTGTATCCATGTGTGGTGGGCTCCTGATCATTCATTTATGCTCAGCGCTGTGAGAAACGGGCCTGAGAATCAGATGCTCTGCT>1034
    201>gtgaaagctctctgcacctgtatccatgtgtggtgggctcctgatcattcatatatgctcagcgctgtgagaaacgggcctgagaATCAGATGCTCTGCT>300

             *         *         *         *         *         *         *         *         *         *
   1035>CTCTGGATCAGCGTGACCTCAGAAATGAAGACGAGAGTCACAAGGCCGCGGCCGCCCGAGGGCAGAGACGTTTACGAGAGCGCAGAGACACACAGAAAGA>1134
    301>CTCTGgatcagcgtgacctcagaaatgaagacgagagtcacaaggccgcggc cgcccgagggcagagacgtttacgagagcgcagagacacacagaaaga>400

             *         *         *         *         *         *         *         *         *         *
   1135>GCCTTTTGTCGTCTGAATTCTTCCGCATCATTTTCTGCATTTCATCAGCTGTGAGTCAAACTCACCCACAAATCTCATTCCTGCCCGCTCTGTGTTTATC>1234
    401>gccttttgtcgtctgaattcttccgcatcattttctgcatttcatcagctgtgagtcaaactcacccacaaatctcattcctgcccgctctgtgtttatc>500

             *         *         *         *         *         *         *         *         *         *
   1235>ACATCACTCACACACACACACAACTGCTCAAATACTGATAAAAACACACACACACACACACACTCATAAATACACACACAAGAACTCAAAAATGCTCA>1334
    501>acatcactcac--acacacacaactgctcaaatactgataaaaacacacacacac--------tcataaatacacacaca------------------>572

             *         *         *         *         *         *         *         *         *         *
   1335>AACACTGATAAACACGCACACACATTCTTAAACACACATACTGATAAACACACactcataaacatgcacaaacacacacaaacactgataaacacacata>1434
    573>------------cacacacacaca----aaaacagaca-actg-----------ct----------------------caaatactg------------->609

             *         *         *         *         *         *         *         *         *
   1435>ttctcataaacacacacacaagcacacacactttctcgttaacatttaggaacaaacactgatacacactgacacacaagtaatcaaaaccgcac>1529
    610>-----ataaacacacacacacacacacacact----------cat-----aa---------atac------acacacaagtaatcaaaaccgcac>669
```

**Figure S3: *peak5* cloned nucleotide sequence**

Zebrafish wild-type reference sequence (795 bp – top line) aligned to cloned *peak5* sequence (669 bp – bottom line). Capitalized nucleotide sequence in top reference sequence indicated region annotated as a peak by ATAC-Seq.

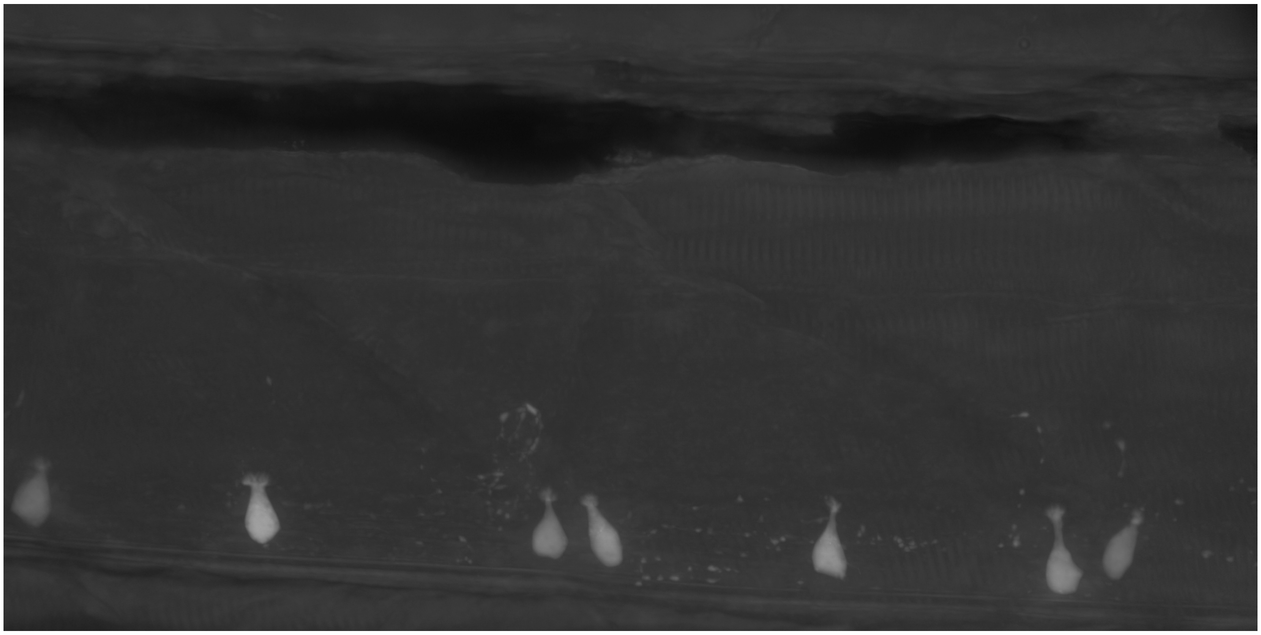

**Figure S4:** ***peak5*** **labels a subset of ventral Kolmer-Agduhr neurons in the spinal cord**

Maximum projection confocal image of *peak5:betaglobin:EGFP* labeled KA neurons in a stable transgenic line.

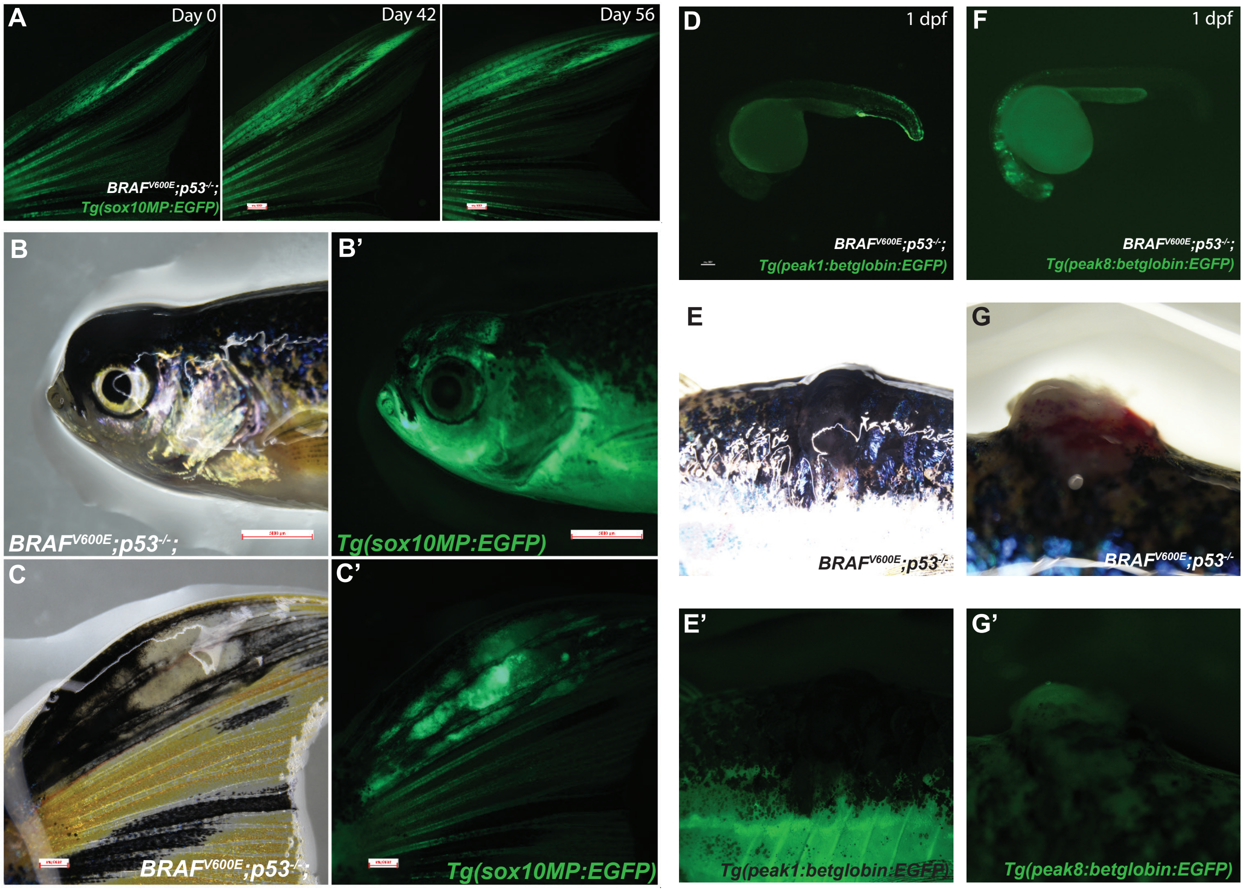


**Figure S5: The *sox10* minimal promoter is active in melanoma. *peak1* and *peak8* are not active in melanoma**

**A)** The *sox10* minimal promoter (MP) is active in melanoma precursor lesions. **B-B')** The *sox10* minimal promoter is active in melanoma tumors on the head and **C-C')** on the tail. **D)** One *peak1* stable line is active embryonically in the fin mesenchyme. **E-E')** *peak1* is not active in melanoma tumors. **F)** A representative *peak8* stable line is active in the CNS embryonically. **G-G')** *peak8* is not active in nearly all melanoma tumors.

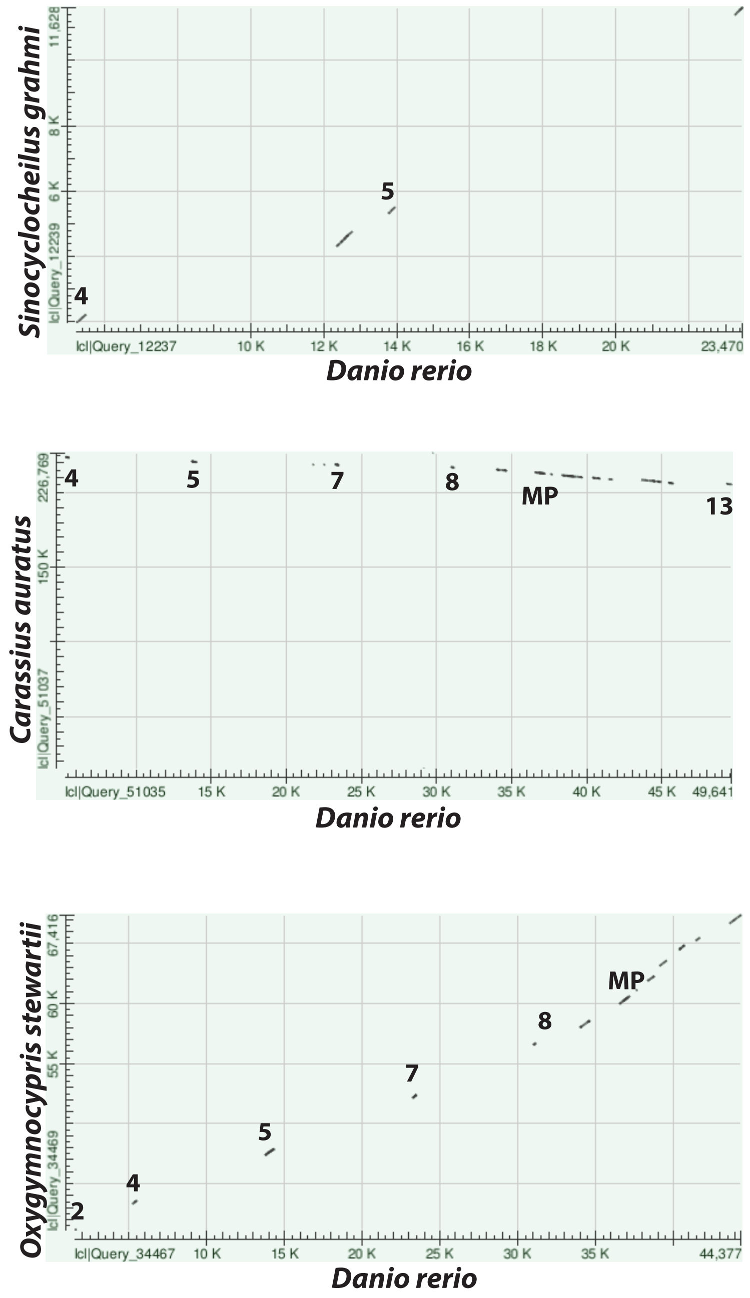


**Figure S6: Conservation of *sox10* enhancers across members of the *Cyprinidae* family**

Dot-matrix view of the alignments of scaffolds containing *peak5* conservation compared to the same *sox10* genomic locus in zebrafish. Numbers indicate regions where peak sequences are conserved. MP = minimal promoter.

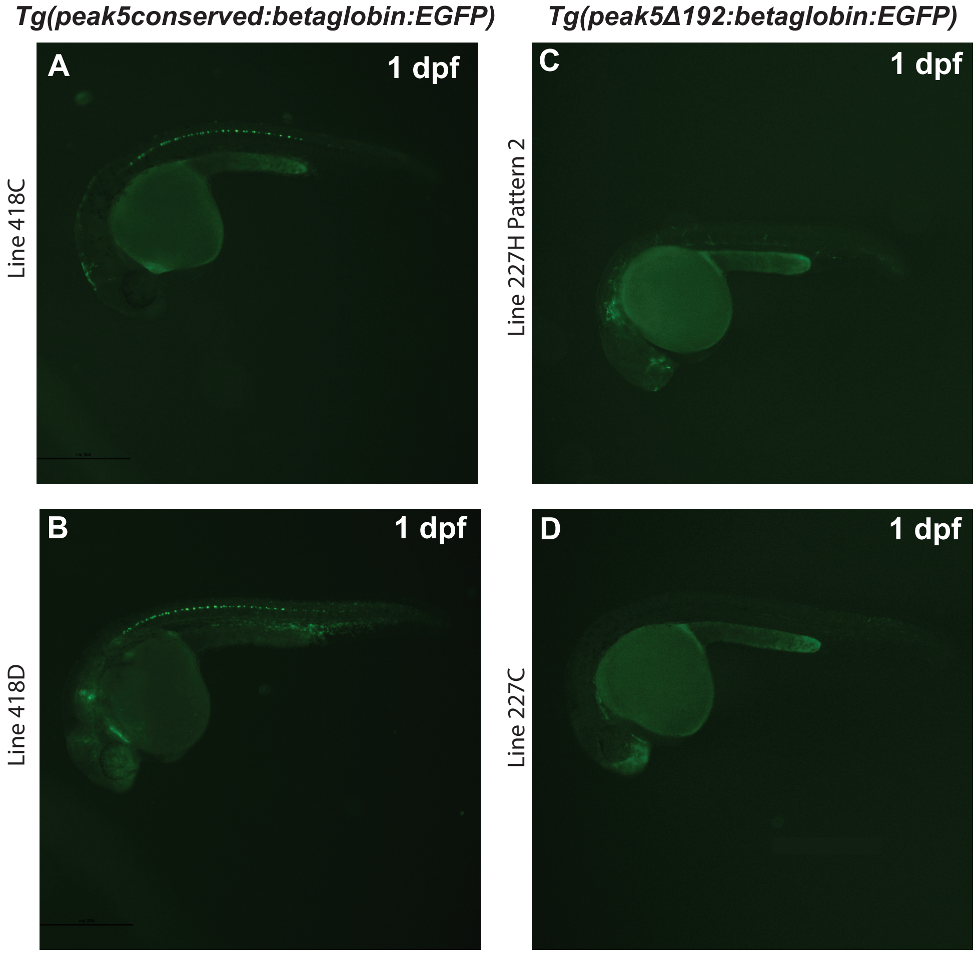


**Figure S7: Additional *Tg(peak_conserved:EGFP)* and *Tg(peak5Δ192:EGFP)* stable lines**

**A and B)** *Tg(peak5Δ192:EGFP)* stable lines exhibit robust *EGFP* expression in KA neurons at 1 dpf. **C)** A second pattern of expression identified from a single founder for *Tg(peak5_conserved:EGFP)* Line 227H. **D)** An additional *Tg(peak5_conserved:EGFP)* stable line exhibits EGFP localization in posterior dorsal NCCs, but does not exhibit any EGFP localization in KA neurons.

### Tables

| Peak Name | Annotated Location | Amplified Location | Distance to TSS (Amplified sequence) |
|---|---|---|---|
| **PEAK 1** | Chr3: 1976452-1976658 | Chr3: 1976387-1976752 | -36.4 KB |
| **PEAK 2** | Chr3: 1977838-1978112 | Chr3: 1977785-1978218 | -35 KB |
| **PEAK 3** | Chr3: 1978414-1978643 | Chr3: 1978343-1978649 | 34.5 KB |
| **PEAK 4** | Chr3: 1981603-1982037 | Chr3: 1981487-1982249 | -31.1 KB |
| **PEAK 5** | Chr3: 1990057-1990509 | Chr3: 1989857-1990651 | -22.8 KB |
| **PEAK 7** | Chr3: 1999674-2000029 | Chr3: 1999609-2000086 | -13.2 KB |
| **PEAK 8** | Chr3: 2007199-2007580 | Chr3: 2006988-2007878 | -5.6 KB |
| **PEAK 11** | -- | Chr3: 2017817-2018186 | 4.9 KB |
| **PEAK 12** | -- | Chr3: 2020003-2020873 | 7.3 KB |
| **PEAK 13** | Chr3: 2025633-2026297 | Chr3: 2025361-2026723 | 12.9 KB |
| **MIN PROMOTER** | Chr3: 2012890-2013261 | Chr3: 2012920-2013269 | 37 BP |
| **NEG CONTROL A** | -- | Chr3: 2002002-2002703 | -10.7 KB |
| **NEG CONTROL B** | -- | Chr3: 1936346-1937052 | -76.3 KB |

**Table 1: ATAC-Seq peak locations in zebrafish (danRer10)**

TSS = Chr3: 2013057. The distance of each amplified peak from the *sox10* TSS was calculated from the middle of each peak.

| Peak Name | | Sequence (5'-3') | Notes |
|---|---|---|---|
| **Peak 1** | | | 376 bp amplified |
| | F | acagtgggaaatgaacagcag | |
| | R | gaagagctgcgttctgctttg | |
| **Peak 2** | | | 434 bp amplified |
| | F | acagttgctatggtaacataacaac | |
| | R | ttgctgaagtggagaaacaac | |
| **Peak 3** | | | 311 bp amplified |
| | F | acaggtttacagaggttaacag | |
| | R | tggaatataaacagaacaaagtcc | |
| **Peak 4** | | | 743 bp amplified |
| | F | tagaaacacaacagaagtgctg | |
| | R | aagccagaatacagcatagc | |
| **Peak 5** | | | 669 bp amplified |
| | F | agtcaatctgacagggtattg | |
| | R | gtgcggttttgattacttgtg | |
| **Peak 7** | | | 478 bp amplified |
| | F | caacaacaatctcagagtgatctc | |
| | R | ttgacttctactgtgtgtgaac | |
| **Peak 8** | | | ~915 bp amplified |
| | F | acatactgtgaacatttcctgaatc | |
| | R | tgtgccatcaattgtgaatcag | |
| **Peak 11** | | | 382 bp amplified |
| | F | cacatgtttcagactgctgaac | |
| | R | ttgatcagttaaatgtgtgtttgag | |
| **Peak 12** | | | 871 bp amplified |
| | F | tgctcaggtcagagtcacagc | |
| | R | tcagtttgtgtcgattgtggtgc | |
| **Peak 13** | | | |
| | F | acttgttaacatgcagctgtca | |
| | R | gttgcgtgagtgtgtacatatg | |
| **Sox10 Minimal Promoter** | | | 351 bp amplified |
| | F | **acaaaaaagcaggctcgcta**cctgtgagaggccaaatattac | bold = Gibson Assembly overlap |
| | R | **cttgctcaccatggtggcga**agtttctccgctagacagtg | bold = Gibson Assembly overlap |
| **Negative Control A** | | | |
| | F | gtacacctgtaatcaaagctgc | |
| | R | gacactgtaatgtacagatgttgc | |
| **Negative Control B** | | | 702 bp amplified |
| | F | tccactgtaagaagcgtaatagg | |
| | R | gaagaagcatagagctgaaagc | |
| **Peak5 Conserved Sequence** | | | 192 bp amplified |
| | F | cagcgtgacctcagaaatgaag | |
| | R | gataaacacagagcgggcag | |
| **Peak 5 Plasmid Mutagenesis** | | **Sequence (5'-3')** | **Notes** |
| **Conserved Deletion** | | | |
| | F | ACATCACTCACACACACAC | |
| | R | ATCCAGAGAGCAGAGCATC | |
| **SoxE Mutations** | | | |
| | F | cct**gcgac**CGTCTGAATTCTTCCGCATC | Bold = SoxE mutations |
| | R | ctc**gccac**GTGTGTCTCTGCGCTCTC | Bold = SoxE mutations |

**Table 2: List of primers used in this study**

| Plasmid Name | Purpose | Source |
|---|---|---|
| **5' Entry Clones** | | |
| p5E*peak1* | Gateway Cloning | Kaufman Lab |
| p5E*peak2* | Gateway Cloning | Kaufman Lab |
| p5E*peak3* | Gateway Cloning | Kaufman Lab |
| p5E*peak4* | Gateway Cloning | Kaufman Lab |
| p5E*peak5* | Gateway Cloning | Kaufman Lab |
| p5E*peak7* | Gateway Cloning | Kaufman Lab |
| p5E*peak8* | Gateway Cloning | Kaufman Lab |
| p5E*peak11* | Gateway Cloning | Kaufman Lab |
| p5E*peak12* | Gateway Cloning | Kaufman Lab |
| p5E*peak13* | Gateway Cloning | Kaufman Lab |
| p5EnegativecontrolA | Gateway Cloning | Kaufman Lab |
| p5EnegativecontrolB | Gateway Cloning | Kaufman Lab |
| p5E*peak5_conserved* | Gateway Cloning | Kaufman Lab |
| p5E-MCS | Gateway Cloning | Tol2 Kit |
| **Middle Entry Clones** | | |
| pME*betaglobin:EGFP* | Gateway Cloning | Tamplin *et al.*, 2011 |
| pENTR-EGFP2 | Gateway Cloning | Addgene 22450 |
| pMEsox10min | Gateway Cloning | Kaufman Lab |
| **3' Entry Clones** | | |
| p3E-polyA | Gateway Cloning | Tol2 Kit |
| **Destination Vectors** | | |
| pDestTol2pA2 | Gateway Cloning | Tol2 Kit |
| **Enhancer Assay Plasmids** | | |
| *peak1:betaglobin:EGFP* | Injection | Kaufman Lab |
| *peak2:betaglobin:EGFP* | Injection | Kaufman Lab |
| *peak3:betaglobin:EGFP* | Injection | Kaufman Lab |
| *peak4:betaglobin:EGFP* | Injection | Kaufman Lab |
| *peak5:betaglobin:EGFP* | Injection | Kaufman Lab |
| *peak7:betaglobin:EGFP* | Injection | Kaufman Lab |
| *peak8:betaglobin:EGFP* | Injection | Kaufman Lab |
| *peak11:betaglobin:EGFP* | Injection | Kaufman Lab |
| *peak12:betaglobin:EGFP* | Injection | Kaufman Lab |
| *peak13:betaglobin:EGFP* | Injection | Kaufman Lab |
| *neg_controlA:betaglobin:EGFP* | Injection | Kaufman Lab |
| *neg_controlB:betaglobin:EGFP* | Injection | Kaufman Lab |
| *sox10min:EGFP* | Injection | Kaufman Lab |
| *peak5Δ192:betaglobin:EGFP* | Injection | Kaufman Lab |
| *peak5_conserved:betaglobin:EGFP* | Injection | Kaufman Lab |
| *peak5_SoxEmut:betaglobin:EGFP* | Injection | Kaufman Lab |

**Table 3: List of plasmids used and/or generated in this study**

| Stable Line | Number of Fish Screened | Number of EGFP+ Tumors | Number of EGFP- Tumors |
|---|---|---|---|
| ***peak5*** **LineA** | 20 | 22 | 2 |
| ***peak5*** **LineB** | 14 | 14 | 1 |
| ***peak5*** **LineC** | 17 | 19 | 1 |
| ***peak5*** **Line129B** | 10 | 12 | 0 |
| **Peak5 Totals** | **61** | **67** | **4** |
| ***peak5Δ192*** **Line328C** | 9 | 0 | 10 |
| ***peak5Δ192*** **Line418C** | 1 | 0 | 1 |
| ***peak5Δ192*** **Line418D** | 12 | 2 | 10 |
| **Peak5Δ192 Totals** | **22** | **2** | **21** |
| ***sox10min*** **LineA1** | 5 | 5 | 0 |
| ***sox10min*** **LineC** | 16 | 15 | 4 |
| ***sox10min*** **Line103B** | 6 | 5 | 2 |
| ***sox10min*** **Line912E** | 7 | 6 | 1 |
| ***sox10min*** **Line1228AC** | 7 | 6 | 1 |
| ***sox10min*** **Line1228A** | 1 | 1 | 0 |
| **sox10 Minimal Promoter Totals** | **42** | **38** | **8** |
| ***peak1*** **LineA** | 6 | 0 | 6 |
| **Peak1 Totals** | **6** | **0** | **6** |
| ***peak8*** **Line918E** | 10 | 0 | 11 |
| ***peak8*** **Line918I** | 9 | 0 | 9 |
| ***peak8*** **Line918F** | 6 | 1 | 5 |
| **Peak8 Totals** | **25** | **1** | **25** |

**Table 4: Numbers of fish screened, and number of EGFP+ and EGFP- tumors for each stable line examined in this study**

| Species | Region of Conservation | Region Used for M-Coffee Alignment |
|---|---|---|
| *Cyrprinus carpio* | scaffold: LG6, chromosome 6:16144792-16145144 | scaffold: LG6, chromosome 6:16144792-16145144 |
| *Sinocyclocheilus grahmi* | scaffold888.1_15: 5319-5517 | scaffold888.1_15: 5319-5517 |
| *Pimephales promelas* | scaffold-281591: 449-637 | scaffold-281591: 449-637 |
| *Carassius auratus* | tig00035131: 220509-220798 | tig00035131: 220509-220757 |
| *Oxygymnocypris stewartii* | isolate Jianluoli-Novo-2018 ctg5171: 47377-47782 | isolate Jianluoli-Novo-2018 ctg5171: 47377 - 47571 |

**Table 5: *peak5* conservation coordinates across members of the *Cyprinidae* family**
